# Cellular and transcriptional trajectories of neural fate specification in sea anemone uncover two modes of adult neurogenesis

**DOI:** 10.1101/2025.06.06.655355

**Authors:** Flora Plessier, Heather Marlow

## Abstract

While neurogenesis is largely restricted to early life stages in animals, some taxa (including cnidarians, planarians and acoels) display lifelong neurogenic abilities. The cellular lineages and transcriptional programs underlying this process remain poorly understood in cnidarians. Combining reporter tracing and single-cell transcriptomics, we identify adult neurogenic trajectories in the sea anemone *Nematostella vectensis*. We uncover two distinct mechanisms: direct differentiation of peptidergic neurons from a multipotent progenitor pool, with identities specified proportionally to existing populations, and a stepwise maturation of cnidocytes (specialized cnidarian neural cells), marked by a transcriptionally distinct intermediate stage. Neural fate commitment is characterized by transient *SoxC* expression, with a homeodomain code associated with neural identities. These modular strategies support continuous neurogenesis and suggest that neural fate specification may use ancestral principles shared with bilaterians. Our study provides a foundational framework for understanding the mechanisms underlying the adult specification of neural cells and their evolutionary diversification across animal lineages.

## Introduction

Neurons are the fundamental units of nervous systems across animals. In most bilaterians (e.g. humans, mice, and flies), neural development is predominantly restricted to early life stages, with limited renewal capacities in adulthood. In contrast, lineages with extensive regenerative abilities and less complex nervous systems, such as cnidarians, acoels, and planarians, maintain ongoing neuronal renewal throughout life^1^. Cnidarians (e.g., sea anemones, corals, hydra, jellyfish), the outgroup to all Bilateria, lack a centralized nervous system, with neurons embedded in diffuse nerve nets^2,3^. However, cnidarian and bilaterian neurons share key effectors, including conserved neuropeptide processing machinery, ion channels, and synaptic proteins, as well as early neurogenic transcription factors like proneural basic helix-loop-helix (bHLH) and SoxB genes, pointing to a common evolutionary origin^4–9^. Major diversifications of transcription factor (TF) families central to bilaterian neural programs -such as Homeodomain, Sox, and bHLH-occurred early in animal evolution, with substantial cnidarian-bilaterian shared expansions, which may have contributed to the diversification of neural identities across both groups^10–12^. Thus, investigating the cell lineages and transcriptional programs involved in neurogenesis in adult cnidarians could provide valuable insights into the mechanisms of lifelong generation of molecularly diverse neural cells across animals.

Cnidarian neural cells encompass ganglion neurons, sensory cells and lineage-restricted cnidocytes (mechano-sensory neural cells). In the genetically tractable sea anemone *Nematostella vectensis,* over twenty subtypes have been molecularly characterized^2,4,13–19^, and its nervous system shows localized condensation of neurite tracts along the primary body axis, indicating a degree of morphological organization^2,20^. Neurons primarily employ a specialized neuropeptide repertoire^4,14–17,19^, along with other neurotransmitters (e.g., glutamate, GABA, but not monoamines)^21–23^. Akin to other highly regenerative species such as planarians, acoels, and annelids^24,25^, cnidarians like *Nematostella* exhibit feeding-dependent indeterminate growth and shrinkage, as well as asexual reproduction^26,27^. Notably, a population of peptidergic neurons (*GLWa*+) scales with body size^14,28^, suggesting that extensive adult neurogenesis occurs. However, the progenitor cell population(s) which enable adult neurogenesis and the transcriptional trajectories underlying this adult program of neural cell fate specification remain poorly understood.

Adult generation of neurons can occur via two distinct yet non-mutually exclusive mechanisms. Each of these mechanisms is associated with different degrees of fate restriction and nervous system organization. In the first, associated with highly organized centralized nervous systems, as in the rodent olfactory system or the zebrafish telencephalon, pools of lineage-restricted neural stem cells maintained within spatially delineated niches can generate a limited variety of neural subtypes that are integrated into nearby structures^29,30^. In the second mechanism, associated with more diffuse centralized nervous systems and nerve nets, as in planarians, acoels and hydra, multipotent migratory stem cells can produce diverse somatic lineages, including most to all neuronal subtypes, under both homeostatic and injury-induced conditions^31–33^. In the adult sea anemone *Nematostella vectensis*, mesenterial *Piwi1*+ cells can give rise to cells expressing the conserved pan-neuronal marker *Elav*^34^, suggesting that they are a source for new neurons. It remains unclear how these putative adult stem cells contribute to adult neurogenesis in cnidarians, including which neurons they give rise to, whether intermediate lineage restriction occurs and what molecular mechanisms accompany fate restriction.

In animals with continuous neural cell renewal -like cnidarians or planarians-, the simultaneous presence of all cellular states (from stem cells to intermediate and mature neurons) presents a significant challenge for identifying cells along one or multiple lineage trajectories. This inability to distinguish temporal intermediates precludes trajectory inference and examination of their transcriptional maturation^35^. Single cell transcriptomics (scRNA-seq) can generate unbiased static snapshots of gene expression profiles across entire nervous systems but lineage inference to investigate cell fate acquisition remains challenging^36^. Imaging of genetic drivers can capture cell dynamics through reporter inheritance, but reconstructing complex trajectories requires multiple targeted genetic tools. These issues are compounded in emerging model systems with limited genetic tools and sparse *a priori* knowledge needed for lineage reconstruction.

To address these challenges, we integrated temporal reporter tracing and single-cell transcriptome profiling to map the lineages and transcriptional programs underlying adult neurogenesis in the sea anemone *Nematostella vectensis*. We find that distributed neurogenesis scales the nerve net with peptidergic neurons generated directly from non-neural-exclusive progenitors, and cnidocytes maturing through a transcriptional intermediate stage before initiating their secondary neural program. These parallel transcriptional lineage strategies employ bilaterian-shared transcription factor classes (bHLH, homeodomain) across neural identities, suggesting common neural development principles for the generation of neural diversity.

## Results

### Distributed peptidergic neuron generation underlies adult nerve net expansion

To support indeterminate growth, an adult neurogenic program involving continuous generation and integration of neurons into the nerve net would be required. To identify such adult-born neural cells *in vivo*, we employed a characterized *Elav* regulatory sequence active in neurons from embryonic to adult stages to drive the expression of a photoconvertible KikumeGreenRed (KikGR) fluorescent reporter^20,37^. We labeled the standing population of *Elav*+ cells by photoconverting this reporter (day 0) and analyzed the body wall nerve net after 4 days. New *Elav*^+^ neurons arising from non-expressing cells would lack the photoconverted tracer and display new reporter expression (hereafter referred to as green cells) (Fig. 1A and Supplementary Fig. 1). Thus, discriminating green neurons from previously generated stably-expressing neurons (red and green co-expressing, orange cells hereafter) or cells that have ceased expressing the reporter (red cells hereafter), allowed us to investigate the dynamics of neurogenesis *in situ*.

**Figure 1.**
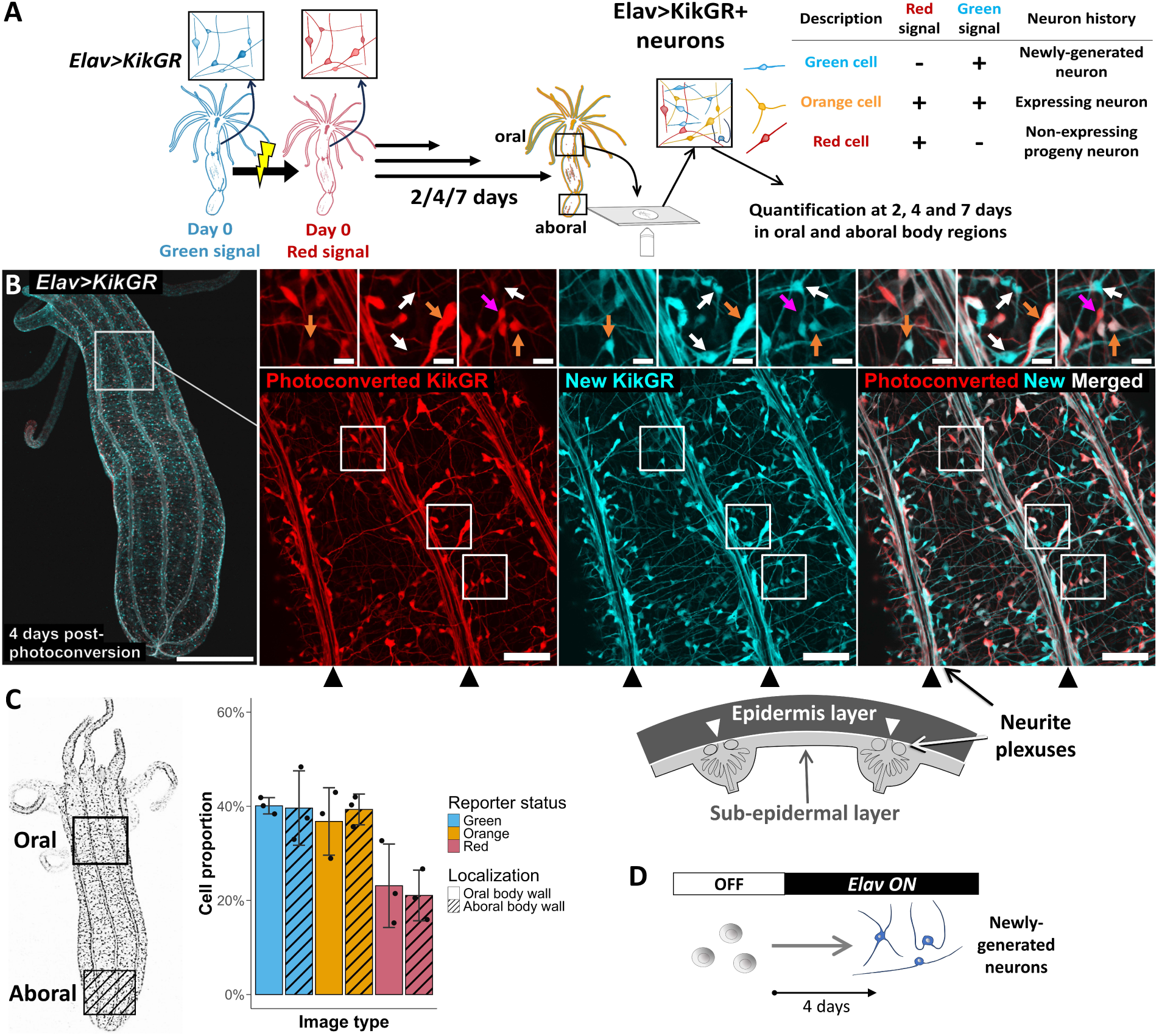
Adult neurogenesis across the primary body axis underlies nerve net expansion. **A** Active adult neurogenesis highlighted by *Elav>KikGR* reporter photoconversion and *in situ* imaging (see Supplementary Fig. 1). At 2,4 and 7 days post-photoconversion, neural cells with the new green reporter signal and lacking the day 0 photoconverted reporter signal are newly-expressing-only cells (green cells, represented in blue throughout the figures). Cells with the photoconverted tracer either also display new KikGR expression (ongoing expression, orange cells) or lack new KikGR expression (progeny cells, red cells). **B** *Elav>KikGR*+ body wall nerve net at 4 days post-photoconversion with 5X global field of view (**left**) and 40X single-cell zoom-in (**right**) images. Photoconverted KikGR day 0 tracer signal is shown in red, and new KikGR expression in cyan. Insets highlight green neurons (white arrows), orange neurons (orange arrows) and red neurons (magenta arrows). Scale bars are 500 µm (left), 50 µm (right) and 10 µm (insets). **C** Proportion of green, orange and red *Elav>KikGR*+ cells detected across raw images in the oral and aboral regions (with hachure pattern) at 4 days post-photoconversion. For each image, cell counts are normalized by the total number of *Elav>KikGR*+ cells detected (n=3 animals, mean ±S.D.). **D** Adult generation of *Elav*+ neurons from *Elav*-negative cells across the body wall nerve net. Data is provided as Source Data File 1.

Imaging *Elav>KikGR*+ animals 4 days post-photoconversion revealed green neurons interspersed with orange neurons in the body wall nerve net (Fig. 1B), indicating ongoing neurogenesis contributing new neurons to the nerve net. Newly-born neurons are dispersed throughout the epidermal and sub-epidermal layers, including in the neurite tracts at the base of mesenteries (Supplementary Videos 1-2). The majority of detected cells (approximately 80%) are green or orange (Fig. 1C), with a small fraction red (Fig. 1B-C); the latter category suggests that a subset of differentiated neurons stop expressing the *Elav* driver. Overall, the large fraction of green cells detected 4 days post-photoconversion is consistent with active neurogenesis contributing new neurons to the body wall nerve net through the rapid generation of *Elav*+ neurons from *Elav*^neg^ cells (Fig. 1D).

To evaluate whether newly generated neurons are exclusively incorporated into specific regions of the nerve net, which could point to a spatially restricted growth zone on the oral side of the primary body axis, we examined both the oral and aboral regions. Green neurons are found in both regions at 2, 4 and 7 days post-photoconversion with similar proportions in each region (Fig. 1C and Supplementary Fig. 2). The synchronous presence of green cells in both regions, coupled with their high fraction, suggest that nerve net expansion occurs through highly active spatially distributed adult neurogenesis, with *Elav*-negative cells differentiating into *Elav*+ neurons within one week.

*Elav*+ neurons encompass a diversity of neural cell states^4^. To investigate the molecular identity of the newly-generated *Elav*+ neural cells, we combined our photoconversion-based temporal recording with single cell RNA-Sequencing (scRNA-Seq) of green, orange and red *Elav>KikGR*+ cells (Fig. 2A and Supplementary Fig. 3/4). We identified 1335 peptidergic neurons from the 3006 *Elav>KikGR*+ sorted cells, highlighted by their shared expression of neuropeptide processing genes (prohormone convertases, amidating enzymes). These encompass over a dozen molecularly distinct subtypes, delineated by their restricted expression of neuropeptide precursors and neuropeptide G-protein coupled receptors (GPCR) (Fig. 2B-C and Supplementary Fig. 5)^15,17^. Notably, we recover all previously described neural populations in the body wall, including a *Prdm14d*+ population (Neural 0)^13–15,17^. As expected, we do not recover *FoxQ2d*+ putative sensory cells and mature cnidocytes, which are not labeled by the *Elav* driver^38^. At 7 days post-photoconversion, we find a majority of the *Elav>KikGR*+ neural cells to be newly-generated green cell (58%), consistent with our *in situ* nerve net imaging (Supplementary Fig. 6). Green cells are distributed among all peptidergic neuronal subtypes (Fig. 2D). Specifically, the number of green cells relative to the combined number of orange and red cells for each cluster was strongly correlated (R^2^=0.96, Pearson’s ρ). This suggests that *Elav*+ neuron subtypes, from the rarer to the more abundant, are generated concomitantly *in vivo*, at a frequency matching their overall proportion in the animal (Fig 2E).

**Figure 2.**
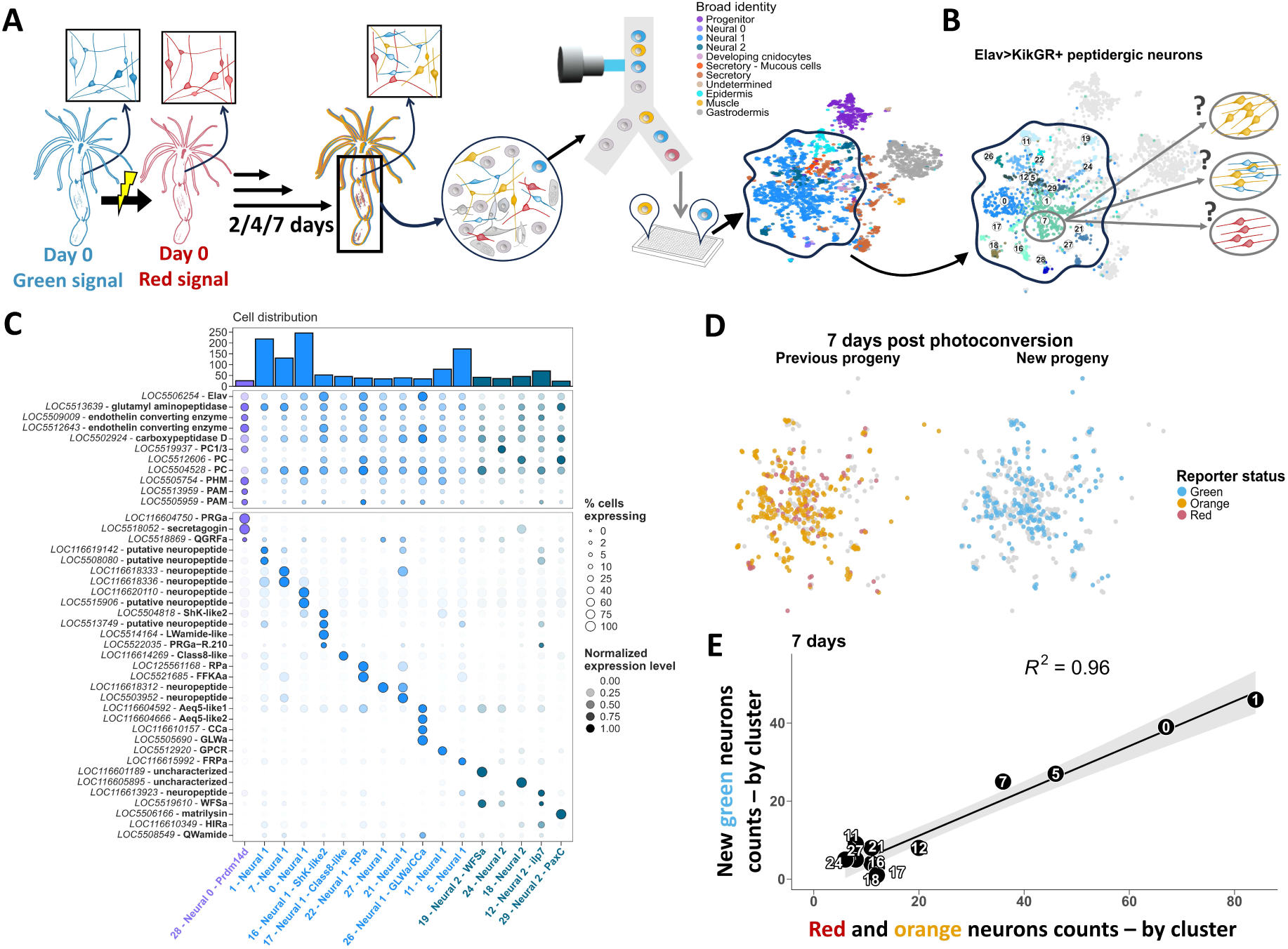
Adult neurogenesis generates all *Elav*+ peptidergic neuron subtypes. **A** Molecular identification of newly-generated *Elav*+ neurons. Plate-based scRNA-Seq (MARS-Seq protocol) allows recording of reporter status post-photoconversion (green, orange or red cell) by flow cytometry for each cell sampled from *Elav>KikGR*+ adult body regions (body wall with mesenteries) at 2,4 and 7 days post-photoconversion. scRNA-Seq transcriptional profiles are then matched to each cell’s reporter status color (Supplementary Fig. 3 and 4). **B** UMAP projection of the 1335 peptidergic neurons from the 3006 *Elav>KikGR*+ sampled cells from 2, 4 and 7 days post-photoconversion with their cluster identifier. **C** Scaled neural marker gene expression across *Elav>KikGR*+ peptidergic neuron clusters. Broadly expressed neural markers and neuropeptide processing genes (**top**) and specifically expressed neuropeptide precursor genes from ^15,17^(**bottom**). Dot size represents the percentage of cells in this cluster which express the gene. Neural cluster identifiers as in **B**. See Supplementary Fig. 5 for expression across all *Elav>KikGR*+ cells. **D** *Elav*^+^ peptidergic neuron scRNA-seq UMAP projection highlighting the overlapping identity distribution between the previously-labeled peptidergic neurons (orange and red cells) (**left**), and the newly-expressing neurons (green) (**right**) at 7 days post-photoconversion. **E** Cell counts by reporter status (green vs. red and orange cells) across peptidergic cell clusters in 7 days photoconverted *Elav>KikGR*+ animals showing a Pearson’s correlation R^2^ of 0.95. Cluster identifiers as in **B-C**. Data is provided as Source Data File 2.

### Progeny of *FoxL2*+ and *SoxC*+ cells encompass neurons, cnidocytes and secretory cells

*In silico* trajectory inference approaches suggest that in the adult, peptidergic neurons, secretory cells, and cnidocytes (cnidarian-specific neural effector cells) are all part of a shared lineage^39^. However, it is unknown whether these populations derive from a common progenitor pool and what molecular features are associated with neural lineage restriction during adult growth. To better understand neural cell fate acquisition, we generated photoconvertible reporter lines for *SoxC* and *FoxL2,* two transcription factors expressed in putative progenitor cells in adults. Earlier work suggests *SoxC* plays a role in the differentiation of cnidocytes, secretory cells and neurons during embryogenesis, and *SoxC* reporter expression has been noted across these cells in the adult^39^. *FoxL2* is a forkhead box transcription factor highly expressed in progenitor cells and tentacular cnidocytes and lowly-expressed in some neurons in *Nematostella*, and in ectodermal neural precursors and neurons in *Hydra*^4,40^, another cnidarian. Together these two reporters allowed us to probe the trajectories of early undifferentiated cell populations that may give rise to diverse neural cells in the adult.

We first investigated the temporal activity of *SoxC* during adult homeostatic growth. Imaging the body wall reveals *SoxC>KikGR* reporter expression in morphologically diverse cells, including neurons, developing and mature cnidocytes, muscle cells, and secretory cells. To determine whether the neural cells might be part of a *SoxC*+ derived lineage, we assessed the presence of red cells post-photoconversion (Fig. 3A-B and Supplementary Videos 3-4). Thus at 7 days post photoconversion, the presence of both green and red mature cnidocytes and other neurons shows these are derived from *SoxC*+ cells, while secretory cells mostly actively express the driver (orange cells). This suggests that *SoxC*+ progenitor cells transiently express the driver early during differentiation and give rise to *SoxC*-negative derivatives including neurons and cnidocytes. Specifically, red neurons and cnidocytes represent cells generated prior to “day 0” of photoconversion while green cnidocytes and neurons are those that developed within 7 days of the photoconversion “chase”. The recovery of primarily red and green cells at earlier time-points (2 and 4 days post-photoconversion), in similar proportions in oral and aboral regions (Supplementary Fig. 7 and Supplementary Videos 5-7), further supports the conclusion that early expressing *SoxC*^+^ cells give rise to *SoxC*-negative derivatives including neurons and cnidocytes.

**Figure 3.**
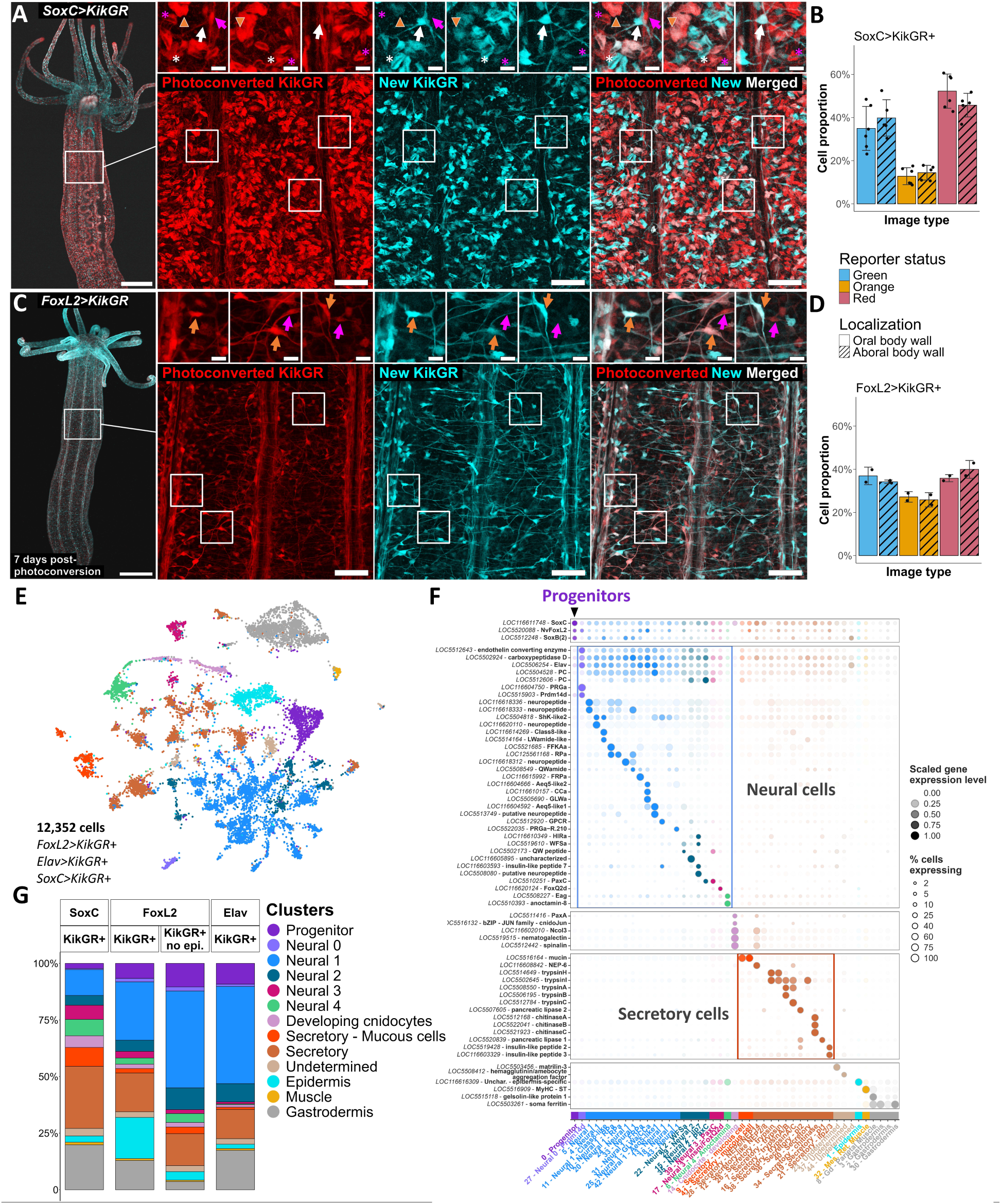
Progeny cells from the *SoxC* and *FoxL2* drivers encompass cnidocyte, secretory and neural cells A,C. *SoxC>KikGR*+ (A) and FoxL2>KikGR+ (C) adult polyp body wall at 7 days post reporter photoconversion. Global field of view (5X, left) and high magnification image (40X extended depth of focus (**A**) or single image (**C**), right) with photoconverted tracer in red and new signal in cyan. *SoxC* driver insets highlight cnidocytes (asterisks) and neurons (arrows) with either only the photoconverted signal (in magenta), or only the new expression (in white), as well as secretory cells with both photoconverted and new signal (orange arrowhead). *FoxL2* driver insets highlight neurons with both photoconverted and new signal (orange arrows) and neurons with only photoconverted KikGR signal (magenta arrows). Scale bars are 500 µm (**left**), 50 µm (**right**), and 10 µm in the insets. **B,D** Proportion of red, orange and green cell bodies detected in the body wall at 7 days post-photoconversion with the *SoxC* (**B**) and *FoxL2* (**D**) drivers. Aboral end body wall images are indicated with a hachure pattern (n=5-6 for the *SoxC* driver and n=2-3 for the *FoxL2* driver, average proportion +/- S.D.). **E** scRNA-Seq UMAP projection of 12,352 *KikGR*+ cells from the *Elav*, *FoxL2* and *SoxC* drivers at 2, 4 and 7 days post-photoconversion, colored by broad cluster identity. **F** Marker gene expression across clusters. For each gene, expression is normalized to the highest expression across clusters. Colors correspond to broad cluster annotations, as in **E**. Dot size represents the proportion of cells expressing the gene in the cluster. Marker genes names are curated from ^13,15,17,38,41,45^, see Supplementary File 1. **G** Cell identity distribution by driver and FACS gating strategy across the *SoxC*, *FoxL2* and *Elav* drivers. For the *FoxL2* driver, the KikGR+ no epi. FACS gate depleted the epithelial epidermis cell fraction. Data is provided as Source Data File 3.

We next analyzed the expression of the *FoxL2* driver. The reporter is expressed in multiple cell types in the body wall, including neurons, mature cnidocytes and epithelial cells, and uncharacterized cells, with sparse secretory and muscle cells (Fig. 3C). We thus investigated whether these differentiated cells derived from a *FoxL2*+ lineage (presence of red cells) or were actively expressing the reporter (orange cells). At 7 days post-photoconversion, we detect diverse red cells, including neurons and mature cnidocytes, suggesting these are non-expressing cells arising from *FoxL2>KikGR*+ cells (Fig. 3C). The sparse green cells present are mature cnidocytes and undetermined cells (Supplementary Fig. 8 and Supplementary Videos 8-9). This shows that *FoxL2^+^* cells give rise to multiple derivatives within 7 days, including neurons. This is consistent with our quantification, which highlights a large fraction of red cells, at all days post-photoconversion assayed and across both sampled regions, showing that this adult developmental process is spatially distributed across the primary body axis (Supplementary Fig. 7). Overall, we uncover neural lineage-derived cells with both the *SoxC* and *FoxL2* drivers, with *FoxL2*+ and *SoxC*+ cells generating *SoxC*^neg^ derivative cells within 7 days, including uncharacterized neurons and mature cnidocytes.

To investigate the molecular signatures of these adult neural developmental lineages, we combined the temporal recording of reporter activity with scRNA-Seq (workflow as in Fig.2A). We sampled *SoxC>KikGR*+ and *FoxL2>KikGR*+ cells at 2,4 and 7 days post-photoconversion (Supplementary Fig. 3/4/9) and linked the molecular identity of each cell to its recorded reporter status (green, orange or red cell). We pooled cells sorted from the *SoxC*, *FoxL2* and *Elav* drivers and annotated 45 clusters using published marker genes^13,15,17,38,39,41–46^(Fig. 3E/F/G). Clusters were then grouped into 13 broad cell classes, with groups Neural 0 to 4 representing *Prdm14d*+ neurons, *Elav*^high^ peptidergic neurons, *Elav*^low^ peptidergic neurons, mixed neural cells including *FoxQ2d*^+^ sensory cells and *Eag*+ mature cnidocytes^47^, respectively. Overall, our cluster resolution is consistent with subtype-specific gene expression patterns within peptidergic neurons, cnidocytes, and secretory cells^17,45^. We recover developing cnidocytes which express cnidarian-specific capsule proteins and known regulators^46,48^, along with mucous cells, and digestive secretory cells (with distinct trypsins, chitinases, and pancreatic lipases)^45^. We identify epidermal epithelial cells, mesenteric retractor and parietal muscle cells^41,44^, cnidarian-specific gastrodermis cells, undetermined cells, and putative progenitor cells (*SoxB(2)a*^high^/*SoxC*^high^). Peptidergic neurons are recovered from all three drivers (*Elav*, *FoxL2* and *SoxC*), while cnidocytes, mucous secretory cells and epithelial epidermis cells are absent from the *Elav* driver (Fig. 3G), consistent with the *in situ* imaging. The partially overlapping molecular identities of these neural populations suggest that peptidergic neurons represent a more restricted lineage distinct from other neural fates.

### *Elav^low^ Piwi1*^+^ progenitors are differentiation-committed

As putative progenitor cells display the highest *SoxC* levels (Fig. 3F/G), and neural cells can arise from unknown *SoxC*+ cells (Fig. 3A), we examined these undifferentiated “progenitor” cells recovered from all drivers as a source population for neural cell derivatives. The progenitor cluster specifically expresses genes associated with proliferative pluripotent cells across metazoans, such as transcription factors from the bHLH (including *Myc*, *atonal*-related, and *achaete-scute*-related genes), Sox B and C families, RNA helicases (*PL10*, *vasa1* and *vasa2*), and RNA-associated proteins (*nanos1*, *nanos2*, *piwi1*, *piwi2*) (Fig. 4A)^32^. Progenitors also display the strongest enrichment of both S-phase and G2/M cycling modules (Fig. 4A). In contrast to progenitors, neurons, cnidocytes and secretory cells have no cell cycle module enrichment, suggesting they are primarily post-mitotic. Epidermis epithelial cells also display a cell cycle module enrichment, consistent with cell division in a subset of epidermis cells^34,49,50^.

**Figure 4:**
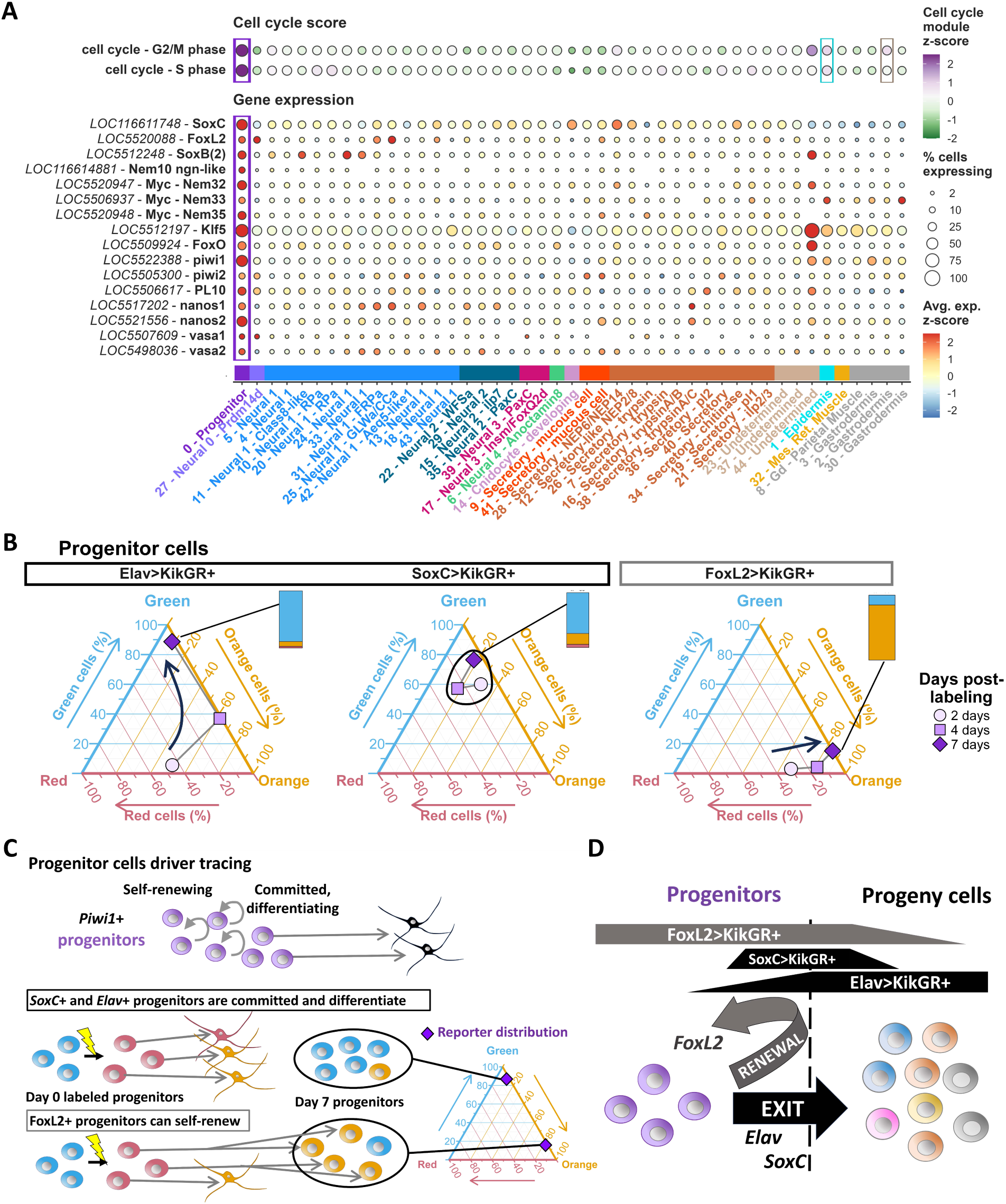
A molecularly homogeneous pool of progenitor cells encompasses both self-renewing and differentiating cells. **A** Molecular profiling of the progenitor cell pool recovered from the *Elav*, *FoxL2* and *SoxC drivers*. (**top**) S-phase and G2/M cycling genes module score across all clusters (same clustering and cluster colors as in Fig. 3E-G). (**bottom**) Gene expression across all clusters. Dot size represents the proportion of cells in each cluster expressing the gene. **B** Diverging temporal fates of progenitors. Ternary diagrams showing the proportion of red, orange and green progenitor cells from the *Elav*, *SoxC* and *FoxL2* drivers at 2,4 and 7 days post-photoconversion, with the same data as a stacked barplot for the 7 day timepoint (all days are also shown as barplots in Supplementary Fig. 11, with the recovered cell numbers indicated). Green cells are represented in blue. **C** Schematics of *Piwi1*^+^ progenitors giving rise to differentiated cells (**top**), with two possible expression dynamics: either expression of the driver occurs only in committed, differentiation-bound progenitors (**middle**) or already in self-renewing progenitors (**bottom**). For both scenarios, the corresponding progenitor cell color proportions at 7 days post-photoconversion are indicated on the right, with a majority of orange progenitor cells at 7 days indicative of driver expression in self-renewing progenitor cells, while a majority of green progenitor cells at 7 days is indicative of driver expression in differentiating and exit-committed progenitor cells. **D** Integrated model of progenitor cell temporal dynamics. Progenitors are a stable, self-renewing pool that express the *FoxL2* driver; initiation of *SoxC* or *Elav* driver expression in progenitors is tied to differentiation initiation within days *in vivo*. Data is provided as Source Data File 4.

Recent work on *Nematostella* stem cells showed that *Piwi1*^+^ dividing cells in the mesenteries can differentiate into *Piwi1*^neg^ *Elav*+ neural cells in the body wall^34^. Here, we show that *Piwi1*+ progenitor cells are also recovered from the broadly-neuronal *Elav* driver^20^ (Fig. 3G). As *Elav* is highly expressed in peptidergic neurons and at low levels in progenitors (Fig. 3F), we hypothesized that the *Elav* positive progenitors identified here are early differentiation-committed cells. If so, progenitor cells recovered at 7 days post-photoconversion should be green, as the labeled progenitors present at day 0 would have differentiated (Fig. 4B-C). We thus evaluated whether the red and orange cell fractions were depleted in 7 day-post-photoconversion *Elav>KikGR*+ progenitors. We find that the majority of progenitors at 7 days (88%) are green. Accordingly, earlier timepoints (2 and 4 days) have intermediate red and orange progenitor cells fractions (Fig. 4B and Supplementary Fig. 10). This is consistent with a scenario in which *Elav*+ progenitors are differentiation-committed exit-bound cells (Fig. 4D).

### *SoxC* is associated with *Piwi1*^+^ progenitor commitment and differentiation

Progenitor cells express *SoxC* and *FoxL2* (Fig. 3F). Our detection of neural derivatives with overlapping class identities from both *SoxC*+ and *FoxL2*+ cells (Fig. 3G) suggests a common *SoxC*+/*FoxL2*+ progenitor pool. However, the presence of green neurons labeled by the *SoxC* driver but only orange and red neurons with the *FoxL2* driver at 7 days post-photoconversion (Fig. 3A/C) reveal distinct temporal expression patterns during differentiation. To better understand this neural differentiation process, we evaluated when progenitors express *SoxC* relative to the initiation of differentiation. Specifically, as endogenous *SoxC* is recovered at a high level in progenitor cells (Fig. 3F), we assessed whether progenitors stably express this driver. If this were the case, using our photoconversion-based temporal recording, we would expect these progenitors to be mostly orange cells. Instead, we find that the majority of sampled *SoxC>KikGR*+ progenitor cells are green cells, at 2-, 4-, and 7-day post-photoconversion (Fig. 4B/C). This pattern suggests that *SoxC* driver initiation in progenitors is linked to a rapid differentiation initiation (Fig. 4C/D).

In marked contrast with the differentiation-committed dynamics observed with both the *Elav* and *SoxC* drivers, most *FoxL2>KikGR*+ progenitors retain their photoconverted label and are either orange or red cells (>87% of progenitors) at 2, 4 and 7 days post-photoconversion (Fig. 4B). This suggests a stable self-renewing pool of progenitors where *FoxL2* driver expression does not imply imminent differentiation (Fig. 4C-D), contrary to the dynamics observed in the *SoxC* or *Elav* drivers. This is further supported by the high endogenous expression of *FoxL2* in progenitors (Fig. 3F), and that, even after 7 days post-photoconversion, *FoxL2>KikGR*+ progenitors remain overwhelmingly orange. Thus, the population of progenitor cells display distinct temporal behaviors (either exit and differentiation or self-renewal) as highlighted by the different drivers. However, the self-renewing (“stem”) and exit-committed/differentiating states are not associated with clearly distinct transcriptional signatures at our sampling resolution. Overall, we uncover a pool of progenitor cells sharing a stem cell-like molecular identity, but with distinct temporal fates *in vivo*: *FoxL2* driver expression in progenitors is not coupled to differentiation, suggesting these *Piwi1*+ progenitors are a standing pool of “stem-like” self-renewing cells, while initiation of *SoxC* or *Elav* within this pool is tied to progenitor state exit and differentiation (Fig. 4D).

### Direct differentiation of peptidergic neurons from *SoxC*^+^/*FoxL2*^+^ progenitors

We then examined the adult developmental lineage of peptidergic neurons, the largest neural class, from these *Piwi1*^+^ progenitors. *Elav*^+^ peptidergic neurons are recovered from all three drivers (*Elav*, *SoxC* and *FoxL2)* (Neural 1 in Fig. 3E/G). To assess whether *Piwi1*^+^/*SoxC*^+^/*Elav^low^*progenitors differentiate into peptidergic neurons, we tracked the derivatives of this population using the *FoxL2* driver (Fig. 5A), as *FoxL2>KikGR*+ progenitors carry the photoconverted label even at 7 days post-photoconversion. We found that over 90% of *Elav*^+^ peptidergic neurons recovered from the *FoxL2* driver are either red or orange cells at 2, 4 and 7 days post-photoconversion (Fig. 5B and Supplementary Fig. 11). Thus, the presence of the tracer label in peptidergic neurons with the *FoxL2* driver after 7 days, together with the large fraction of red cells, are consistent with our proposed model of active neuron generation from *FoxL2^+^*/*Piwi1^+^* progenitor cells. As *SoxC* expression in *FoxL2^+^*/*Piwi1^+^* progenitors is transient and linked to differentiation initiation, if these *Piwi1*^+^/*SoxC*^+^ cells develop into *Elav*^+^ peptidergic neurons, we predicted that the peptidergic neurons recovered from the *SoxC* driver should encompass both red cells (cells generated prior to photoconversion), and an increasing fraction of orange and green cells (newly generated cells) at later timepoints (Fig. 5A). Therefore, we examined whether we could recover both green, orange and red peptidergic neurons from the *SoxC* driver at all days post-photoconversion sampled (Fig. 5B). Overall, the simultaneous presence of red, orange, and green peptidergic neurons from the *SoxC* driver, consistent with our *in situ* nerve net imaging data (Fig. 3A/C), supports our model of peptidergic neurons differentiating from *FoxL2*+/*SoxC*+ progenitors, with a transient early expression of *SoxC*.

**Figure 5:**
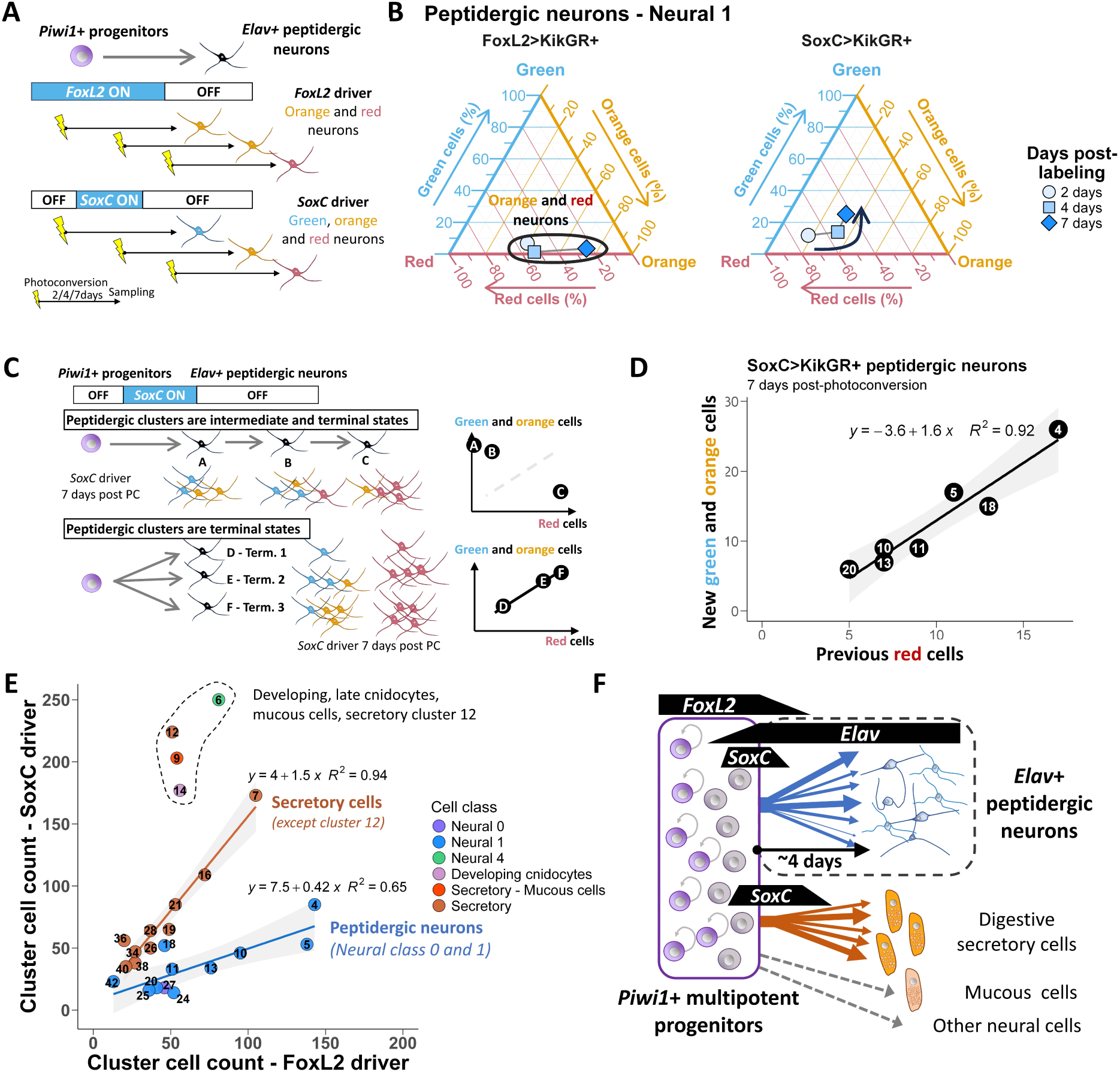
Generation of *Elav*+ peptidergic neurons subtypes from *FoxL2*+/*SoxC*+ progenitors. **A** Putative *SoxC*+/*FoxL2*+ *Piwi1*+ progenitors differentiating into *Elav*+ peptidergic neurons highlights predicted differential neuron reporter status between the *FoxL2* and *SoxC* drivers, with only red and orange neurons expected from the *FoxL2* driver while red, orange and green neurons would be recovered from the *SoxC* driver at 2, 4 and 7 days post reporter photoconversion. Green neurons are shown in blue. **B** Green, orange and red cell proportions for *Elav*+ peptidergic neurons recovered from the *FoxL2* (left) and *SoxC* (right) drivers at 2, 4 and 7 days post-photoconversion. *Elav*^+^ peptidergic neurons from the *FoxL2* driver are red or orange. Green, orange and red *SoxC* driver neurons are recovered, with the arrow indicating the increased fraction of newly generated progeny neurons (green and orange cells) at later timepoints. **C** Schematics for two scenarios of peptidergic neuron development: distinct *Elav*+ neuron clusters represent (top) temporal differentiation intermediates or (bottom) terminally differentiated populations concomitantly generated. As the SoxC driver transiently labels differentiating cells, tracking the ratio between newly arising green+orange neurons and previously-generated red neurons recovered from each peptidergic cluster can discriminate between the two scenarios. This ratio is similar across clusters that are terminal states (bottom) but not for clusters that are transient intermediates (top). Term.: terminally differentiated neural state. Letters indicate neuron clusters. **D** *SoxC>KikGR*+ peptidergic neuron clusters distribution at 7 days post-photoconversion highlights a similar fraction of green and orange cells relative to the red cells across all peptidergic clusters. **E** *Elav*+ peptidergic cluster cell counts recovered from the *SoxC* versus the *FoxL2* driver (*KikGR+* cells, all days post-photoconversion), in blue (R^2^=0.65), with digestive secretory cells in orange, and other neural and secretory cell clusters for comparison. **F** *Elav*^+^ peptidergic neuron development model, with differentiation into distinct peptidergic neural subtypes from peptidergic-committed *Piwi1*^+^ multipotent cells within 4 days in adults. Data is provided as Source Data File 5.

To identify the neural lineage(s) giving rise to peptidergic neuron subtypes, we next investigated whether the distinct *Elav*^+^ peptidergic neurons (Fig. 2C)^14,17^ are terminally differentiated, or whether some are temporal intermediates. We thus compared the distribution of the newly generated cells (green and orange cells from the *SoxC* driver) to the standing cell pool (red cells) across all *Elav*^+^ peptidergic neuron clusters (Fig. 5C and 5D). As the *SoxC* driver is transiently expressed during differentiation, if cell clusters were temporally sequential intermediates, then the ratio of new cells with that identity to the old cells should not correlate, as the previous cells would have matured into a different molecular identity (Fig. 5C – top scenario). At 7 days post-photoconversion, longer than the peptidergic maturation timeline, we found a strong linear relationship between newly and previously generated neurons across peptidergic subtypes (R^2^= 0.92, Fig. 5D). Although the number of sampled cells for the less abundant subtypes limits conclusive findings across all clusters, our analysis of the more abundant clusters suggests that *Elav*^+^ peptidergic neurons are distinct terminal states, rather than intermediate states.

Overall, our data are consistent with closely related peptidergic subtypes being generated from the *Piwi1*+ multipotent population, with their relative ratio fixed at an early peptidergic-committed stage. Furthermore, if there is a shared peptidergic-committed lineage, as the *FoxL2*, and *SoxC* drivers label both the *Piwi1*+ source population and its derivatives, then the relative abundance between peptidergic subtypes would be constrained by this shared lineage and would thus correlate across the *SoxC* and *FoxL2* drivers. Indeed, we find that the relative proportion of peptidergic cells across drivers is correlated and that this relationship is specific to the Neural 1 *Elav*^+^ peptidergic subtypes and the *Prdm14d*^+^ neurons (Neural 0) (Fig. 5E); while cnidocytes and secretory cells do not share the same relationship. This peptidergic lineage restriction within a subset of progenitors is further supported by reanalysis of RNA-Seq data from a *Prdm14d* reporter line, which labels both *Prdm14d*+ neurons (Neural 0 in Fig. 3F), and uncharacterized progenitors at low levels^13^. Genes enriched in *Prdm14d>GFP*+ cells are expressed here in *Prdm14d+* neurons, as well as across non-*Prdm14d*-expressing peptidergic subtypes; and not in secretory cells or other neurons (Supplementary Fig. 12), suggesting that peptidergic-committed *Piwi1*^+^/*Prdm14d*^low^ progenitors are indeed restricted to peptidergic neural fates. Accordingly, genes enriched in *Prdm14d>GFP*-negative cells are expressed across secretory cells and other neural cells, further supporting that these are not part of the shared peptidergic lineage.

Taken together, the data supports a model in which rapid generation of distinct *Elav*^+^ peptidergic neuron subtypes from peptidergic-restricted *Piwi1*^+^/*SoxC*^+^/*Elav^low^*cells underlies homeostatic nerve net scaling (Fig. 5F).

### Neural cells arising from distinct lineages deploy independent TF-effector gene modules

Sea anemone neural cells encompass peptidergic neurons, putative sensory cells, and cnidocytes, cnidarian-restricted mechano-sensory neural cells^2,51^. Having addressed the progression of peptidergic *Elav*^+^ neurons, from *Elav*^low^/*SoxC*^+^/*FoxL2*^+^/*Piwi1*^+^ progenitors to differentiated cells, we examined whether neurons and cnidocytes share a similar differentiation trajectory. Cnidocytes, like peptidergic neurons, develop from *SoxC*+ cells (Supplementary Fig. 13). Thus, only orange and green developing cnidocytes are detected *in situ* and in the scRNA-Seq data with the *SoxC* driver at 2,4 and 7 days post-photoconversion). Mature cnidocytes do not actively express *SoxC* (Fig. 3A/F). Accordingly, we find red mature cnidocytes at 2 days post-photoconversion, with newly generated orange and green mature cnidocytes appearing at 4 and 7 days, respectively (Supplementary Fig. 13 and Supplementary Videos 3-7). Thus, both peptidergic neurons and cnidocytes transiently express *SoxC* during early differentiation. *Elav*+ peptidergic neurons are recovered across all drivers (Neural 0/1/2 in Fig. 3G), but late cnidocytes (Neural 4 class) are not recovered from *Elav* driver-derived sorted cells (Fig. 3G, also in ^20^). Thus, our findings are incompatible with a *SoxC*+ neural-specific bipotent intermediate state.

We then investigated the transcriptional signatures of neural fate acquisition in peptidergic neurons and cnidocytes, two distinct cellular lineages (Fig. 6A). Cnidocytes mature through two sequential transcriptional phases (Supplementary Fig. 13). Initially, during the developing cnidocyte stage, cnidarian-specific genes are used to build their specialized capsule organelle (intracellular harpoon-like structure). Subsequently, developing cnidocytes initiate a neural transcriptional program, which includes the expression of neural effector genes, such as the *Eag* (Ether-à-go-go family) voltage-gated K^+^ channel^47^ and a voltage-dependent L-type calcium channel subunit beta-2 (*Cacnb2*)^51^. This biphasic cnidocyte effector program is characterized by a shift in transcription factors usage. Early in development, cnidocytes express *PaxA*, *cnidoJun*, *NR12*, multiple zinc finger and basic leucine zipper (bZIP) transcription factors, which are associated with capsule formation^42,52,53^. In contrast, the secondary stage is marked by a distinct regulatory signature defined by expression of *nk6*, uncharacterized *Sox* and *Myc* genes and additional bZIP genes (Supplementary Fig. 13), indicative of the onset of the neural program. Thus, cnidocytes undergo two distinct transcriptional phases, with the sequential deployment of a cnidarian-specific capsule program, followed by a neural program. This two-phase transcriptional process is supported by reanalysis of RNA-Seq data from a *PouIV* knockout (KO) mutant^18^, in which immature cnidocytes accumulate. Thus, genes upregulated in *PouIV* KO juvenile polyps correspond to those expressed in developing cnidocytes whereas PouIV KO downregulated genes overlap with those expressed in both developing and late cnidocytes (Supplementary Fig. 14). These findings support a biphasic transcriptional maturation and suggest that when the developing cnidocyte program is blocked, the secondary neural program is not initiated. Thus, this sequential maturation is a two-step process with a clear transcriptional intermediate program that is deployed, then cleared prior to the initiation of the neural program, revealing a different neurogenic strategy from that of peptidergic neurons.

**Figure 6:**
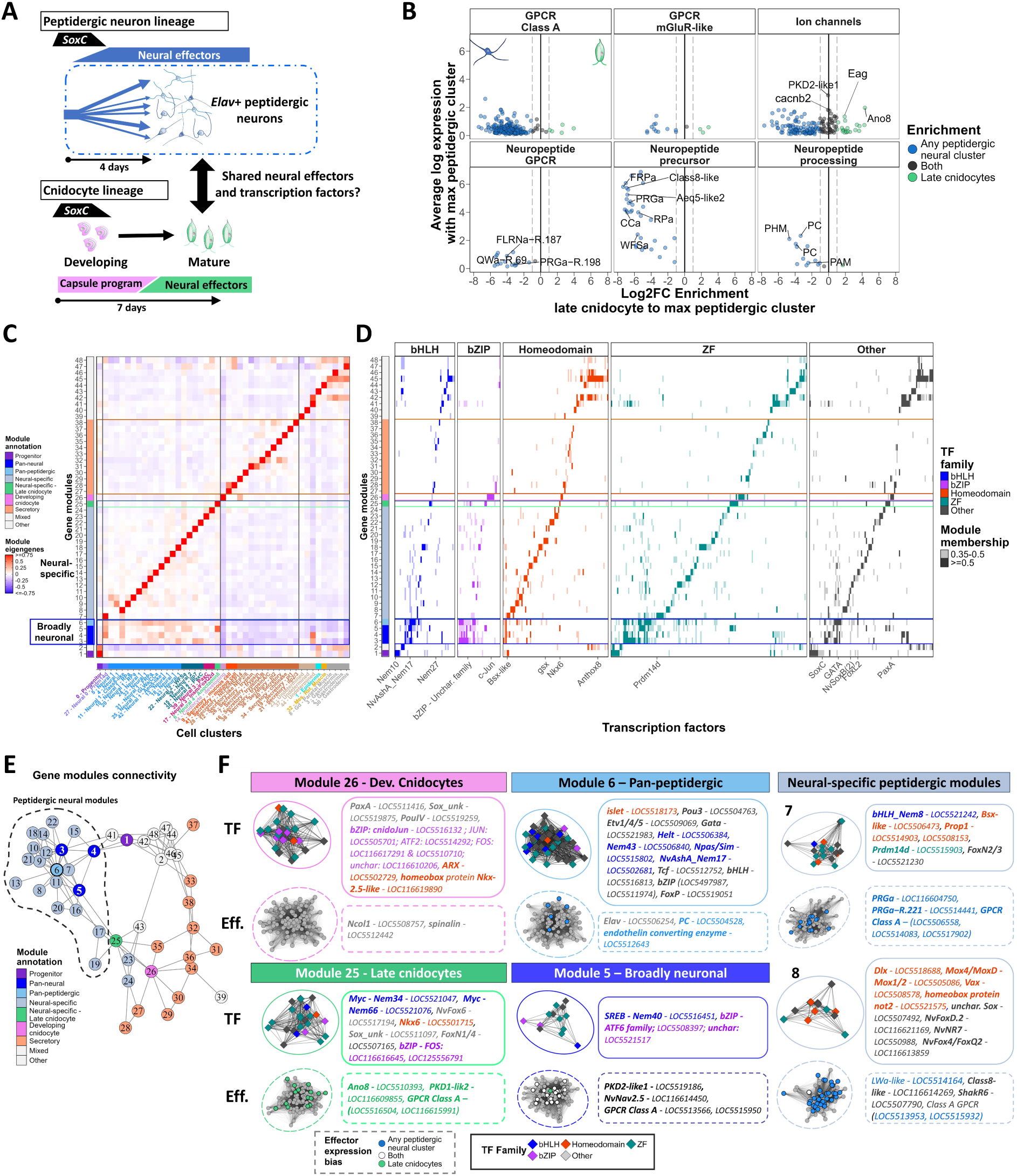
Peptidergic neurons and cnidocytes deploy distinct TF-effector gene modules with bHLH and bZIP factors for broad neural identity and a subtype-specific homeodomain code. **A** Neural cell differentiation in adults. *Elav*+ peptidergic neurons (**top**) and cnidocytes (**bottom**) acquire their mature neural identities through distinct cellular trajectories with an initial transient expression of *SoxC*. **B** Neural effector differential expression between *Elav*+ peptidergic clusters and late cnidocytes. Average expression log2 fold-change enrichment between the late cnidocyte cluster and the maximum expressing peptidergic cluster. Dashed line represents a 2-fold change in expression levels between the peptidergic cluster with the highest expression and the late cnidocyte cluster. Neuropeptide GPCRs and neuropeptide precursors annotated from^15^. **C** WGCNA gene modules eigengene value across cell clusters. Gene modules (rows) are annotated as neural-specific, broadly neural or secretory-specific. Cell clusters (columns) colors as in Fig. 3G. **D** Transcription factor membership across WGCNA modules (same rows as in C) for bHLH, bZIP, homeodomain, zinc finger and other transcription factor families. Each transcription factor is a column, with low (semi-opaque, membership >=0.35) or high (opaque, membership >=0.5) module membership grades to highlight TF associated to multiple modules. Horizontal lines demarcate WGCNA modules associated with broad or restricted neural identities, mature cnidocytes, developing cnidocytes or secretory cells (respectively bottom to top, as in C). **E** Gene modules top co-expression relationships with the gene module identifiers, same module colors as in C. Broadly neural gene modules are in light blue. **F** Neural modules overview. Co-expression networks for broad neuronal and subtype-specific neural gene modules. (**above**) Transcription factors in each module, colored by family. (**below**) Effector genes colored as in B, with neural effectors expressed across both late cnidocytes and peptidergic neurons (in black in B) in white. Pan-peptidergic module 6, broadly neural module 5 and two neural-specific modules (7 and 8) are highlighted, with all neural-associated modules in Supplementary Figure 15. Data is provided as Source Data File 6.

Since mature cnidocytes and peptidergic neurons acquire their neural identities through distinct processes, we further examined whether they converge on a shared neural transcriptional signature (Fig. 6A). We find that the expression profiles of neural effector genes are distinct (Fig. 6B). Neuropeptide precursors, processing enzymes and neuropeptide GPCRs are expressed in peptidergic neurons but not in late cnidocytes, suggesting that peptidergic transmission is not utilized by cnidocytes. Other neural-expressed GPCRs and ion channels are specific to either late cnidocytes or peptidergic neurons, with only a limited overlap between the two. Overall, these results demonstrate that mature cnidocytes and peptidergic neurons have separate neural effector expression profiles.

Despite the distinct differentiation trajectories and effector profiles of these neural cells, we examined whether they shared pan-neural transcription factors. Using weighted gene co-expression network analysis (WGCNA)^54^ to generate modules of genes whose expression is highly correlated, we identify 48 cell-type specific and broader class-specific modules across all cell types (Fig. 6C). Each neural identity combines a cell-type specific module (modules 7-21 and 25) with shared neural module(s) (modules 3-6) (Fig 6C and Supplementary Fig. 15). This is specific to neural cells, and contrasts with the lack of a shared pan-secretory module for the dozen molecularly distinct secretory subtypes. Among the shared neural modules, we first identify a pan-peptidergic module (Module 6) for peptidergic neural identity. This module encompasses pan-peptidergic effectors and transcription factors including a GATA factor, a ETS factor, and bHLH factors such as *NvAshA*, an *achaete-scute*-related gene involved in embryonic peptidergic specification^11,55^ (Fig. 6D and Supplementary Fig. 12). Thus, we propose that this set of transcription factors underlies early peptidergic maturation, consistent with our model of *Elav*-positive peptidergic neurons arising from a shared cellular lineage. Top co-expression relationships between modules further highlight this pan-peptidergic module 6 as a “hub” for peptidergic subtype-specific gene modules 7 to 21 (Fig. 6E). Secondly, we find a smaller pan-neural module partially shared between late cnidocytes and peptidergic neurons (Module 5). It includes few neural effectors and transcription factors, such as the SNAIL family C2H2 zinc finger transcriptional repressor *Scratch*, also used in vertebrate neural maturation^56,57^. While this module spans both late cnidocytes (neural phase) and peptidergic neurons, it is not utilized in the first stage of cnidocyte maturation (capsule phase) and thus does not correspond to a common lineage intermediate. Furthermore, as our early peptidergic neural fate specification model suggests, progenitor Module 1 includes the *atonal-related* bHLH transcription factor *Nem10* (*NvAth-like*/*NvArp3),* an embryonic neurogenesis regulator^11,58,59^ (Fig. 6D). In contrast, the developing cnidocyte module (Module 26) encompasses all known embryonic regulators of early cnidocyte development (*PaxA*, *the bZIP cnidoJun*, *NR12*, and zinc finger factors^18,42,52,53^). While cnidocyte and peptidergic neural programs exhibit limited transcriptional convergence and rely on distinct maturation strategies, there are a few shared factors which suggest some pan-neural shared regulation.

Overall, we find classes of transcription factors systematically associated with pan-(bHLH, bZIP classes) and subtype-specific (Homeodomain, zinc fingers classes) neural modules. bHLH transcription factors play proneural roles in bilaterians, and cnidarian orthologs of the *achaete*/*scute* and *atonal*/*NeuroD* ^51–54,57,58^ are associated here with shared pan-neural modules and broad peptidergic and cnidocyte neural fate, along with *Myc* family genes (Fig. 6D and Supplementary Figure 15). Sets of bZIP factors are enriched in pan-peptidergic, pan-neural, developing and mature cnidocytes modules (respectively modules 6, 3-5, 26 and 25). In addition to class-specific neural bHLH and bZIP factors, each neural subtype is defined by a unique homeodomain and zinc finger code (Fig. 6D and Supplementary Figure 16). Distinct homeodomain factors, with minimal overlap, specify peptidergic neurons, late cnidocytes, and sensory cells; a strategy exclusive to neural cells and absent in secretory cells. This homeodomain code is associated with subtype-specific effectors, including ion channels, neuropeptide precursors, and receptors (Fig. 6D/F). Developing and late cnidocytes also exhibit unique homeodomain and ZF transcription factors, with only *PouIV*, a BTB (Broad complex, Tramtrack, Bric-à-brac family), and an H2C2 ZF transcription factor shared between the two phases.

Together, these analyses highlight that adult neural identity integrates three transcriptional modules: (1) a pan-peptidergic module defining peptidergic neural identity, (2) a small, broadly neural module partially shared between late cnidocytes and peptidergic neurons, and (3) subtype-specific programs with a homeodomain code. Cnidocytes undergo a two-step differentiation process, each phase employing distinct transcription factors separate from those in peptidergic neurons. While peptidergic neurons and cnidocytes follow independent cellular trajectories and transcriptional programs to establish their mature neural identities, they utilize similar transcription factor classes -bZIP and bHLH factors act broadly within neural classes, whereas homeodomain and zinc finger factors are subtype-s pecific. These modular neural renewal strategies are unique to neural cells and are not employed in digestive secretory cells, despite their shared cellular mode of renewal with peptidergic neurons.

## Discussion

A diversity of animals, including cnidarians, planarians and acoels, display indeterminate growth, necessitating lifelong (re)generation of multiple neuron types^1,24,25^. Here, we elucidate the cellular and transcriptional trajectories underlying adult neurogenesis in the sea anemone *Nematostella vectensis*. Using reporter tracing, nerve net imaging and single-cell transcriptomics, we show that active neurogenesis expands the nerve net along the primary body axis through two distinct pathways: (1) peptidergic neurons, the major neural cell class, arise directly from multipotent progenitors, with subtype identities specified in proportion to existing peptidergic populations, and (2) cnidocytes undergo a two-stage maturation process characterized by a transcriptionally distinct intermediate phase, marked by phase-specific transcription factors. In the context of indeterminate growth, our observations show that neurogenesis occurs across the primary body axis, without a restricted growth zone, reminiscent of the distributed processes described in planarians and acoels^1,25^. The high neurogenic rate uncovered -over one peptidergic neuron produced for two existing ones per one-week period-, consistent between unbiased *in situ* imaging and targeted scRNA-Seq data from multiple reporters, aligns with the feeding-modulated exponential growth rates observed in adults^26^. Importantly, the concomitant generation of neural subtypes in fixed proportions occurs independent of injury. This suggests that regenerative capacities may be rooted in continuous cell renewal rather than exclusively as a damage response. The occurrence of homeostatic neural renewal is tightly associated with extensive injury-induced regeneration and asexual reproduction abilities across metazoans^1^. Although it remains unclear whether indeterminate growth could represent an ancestral condition^60^, we propose that the refinement of animal body plans, with matrix-encased compartmentalized organs, may have limited the potential for continuous cell renewal across all cell types, and thus indeterminate growth, asexual reproduction and regenerative abilities^60^.

We show the continuous differentiation of cells from the adult stem-like progenitor compartment^34,61,62^ into multiple cell types, including neural and secretory cells. The transcriptional signature of this progenitor pool resembles that of adult multi-to pluripotent stem cells found in planarians, acoels and in *Hydra* and *Hydractinia*^31,32,63,64^, pointing to shared molecular mechanisms of stem cell potency in animals. Notably, our study demonstrates that cells (*SoxC*^+^ or *Elav*^+^) within this population are fate-committed, and rapidly differentiate, even in the absence of clear transcriptional differences with the rest of the progenitor compartment at our resolution. These findings indicate that differentiation initiation occurs rapidly within the self-renewing progenitor pool, without clear transcriptional intermediates for distinct fates. This suggests that fate decisions are made alongside the cell cycle, as proposed for planarian neoblasts^65^. This contrasts with neural development in fixed-growth animals, where transit-amplifying intermediates, such as *Drosophila* neuroblasts, or mammalian apical radial glia, sequentially generate neurons during embryonic neurogenesis^66^. In *Nematostella*, direct neurogenesis may enable rapid neuron production to sustain indeterminate growth, with all body wall peptidergic neuron subtypes arising from undifferentiated progenitors per four-day period. However, this could constrain molecular diversity, producing fewer neuron types compared to fixed-growth bilaterians.

Our study further uncovers lineage biases within the stem-like progenitor pool. Although all three drivers label the progenitor compartment, their resulting progeny show only partial overlap, with differential neural and secretory cell class recovery. This suggests two possible models for neural cell fate determination. One model proposes a shared “neuroglandular” lineage, where cells exiting the progenitor state are committed to either a peptidergic or digestive secretory cell fate, consistent with previous hypotheses^39^. Alternatively, a second model proposes that commitment to these fates occurs earlier within the progenitor compartment, with distinct cellular lineages forming upon exit. Our data, along with the reanalysis of a *Prdm14d* driver strongly support the latter scenario. This is also consistent with the *insm1* reporter data^67^, as *insm1* is expressed in progenitors, as well as in secretory cells and neurons in adults. Furthermore, the concurrent generation of peptidergic neurons and digestive secretory cells, in proportions that reflect their relative subtype abundance, and the absence of a cycling signature in either population, raises the possibility that subtype identity is pre-determined upon exit from the progenitor state.

In contrast to the direct generation of peptidergic neurons, cnidocytes, the hallmark stinging cell of the phylum, undergo a distinct two-step transcriptional maturation. They transition from a cnidarian-specific capsule-forming program to a bilaterian-like neural program, each associated to distinct sets of transcription factors. The secondary neural phase of cnidocyte development shares some transcription factors with peptidergic neurons, suggesting limited transcriptional convergence across neural classes. Cnidocytes are conserved across all cnidarians^46,68–70^, but their renewal mechanisms vary. In hydrozoans, which diverged from anthozoans over 500 million years ago, such as the jellyfish *Clytia hemispherica*, they arise from stem cell niches in tentacle bulbs, while *Hydra* produces migratory nests of 4/8/16/32 cells from multipotent interstitial stem cells^71,72^. In contrast, *Nematostella* (an anthozoan) shows dispersed early-stage cnidocytes in the epidermis and mesenteries, even as it remains unclear whether they arise from the stem-like progenitor pool (also in^34,62^). This highlights the evolutionary plasticity of cnidocyte development, despite their conserved capsule genes. Thus, investigating whether orthologous transcription factors and neural effectors genes are used in cnidocytes and peptidergic neurons across cnidarians could shed light on neural cell evolution and diversification, particularly considering their lineage-specific neural effectors expansion (e.g., voltage-gated potassium channels, GPCRs^15,73^).

Although cnidocytes and peptidergic neurons use distinct renewal modes, their early development is marked by transient *SoxC* expression, followed by downregulation during maturation. This parallels vertebrate neurogenesis, where SoxC factors (SOX4/11/12) play a critical role in neural progenitor differentiation^74^, underscoring the evolutionary conservation of neurodevelopmental pathways across eumetazoans, despite lineage-specific diversification. Overall, we propose a model where *SoxC*, *SoxB(2)* and distinct bHLH and bZIP transcription factors are associated to neural class specification, while homeodomain and zinc finger genes specify subtype identities. Intriguingly, the homeodomain code delineating neural identities in *Nematostella* is unique to neural cells and absent in digestive secretory cells. This is not clearly associated to axial patterning, as body wall neurons do not appear clearly spatially segregated along the primary (oral-aboral) or secondary (directive) body axes, across currently characterized populations^13,14,17,18,28,38^. Furthermore, the neural-subtype-specific homeodomain proteins are distinct from the ANTP-class Hox genes specifying the secondary (directive) body axis^75,76^, expressed in cnidarian-specific gastrodermal cells in adults. In bilaterians, homeodomain transcription factors are essential for primary body axis (antero-posterior) patterning (ANTP-class Hox genes), and act as terminal selectors of neural identity^77–80^. Thus, given the early diversification of homeodomain family genes before the cnidarian-bilaterian split, their neural subtype-specific expression in *Nematostella* suggests an ancestral role in neural subtype specification, separate from spatial patterning^10,81^. Supporting this hypothesis, the POU-family homeodomain gene *PouIV/Brn3* is critical for cnidocyte maturation^18,82^, as its removal results in blocked immature cnidocytes that fail to initiate their secondary neural program. Systematic functional studies of homeodomain-class genes during adult neurogenesis would help shed light on this evolutionary link.

Our identification of the populations, lineages, and transcriptional programs underlying adult neural cell generation in a sea anemone reveals how indeterminate growth organisms scale their nervous system. We uncover two distinct modes of renewal across the main neural classes, shedding light on the strategies underlying lifelong neurogenesis. The shared molecular features with bilaterians point to deeply conserved neural specification mechanisms, tracing back over 600 million years to the Precambrian. These findings provide new insights into the molecular basis of adult neurogenesis and the evolutionary diversification of neurons across animal lineages.

## Methods

### Transgenic line generation

For the *FoxL2>KikGR*, *SoxC>KikGR* and *Elav>KikGR* reporter line generation, plasmid constructs generated from a pNvT-MHC::mCh vector backbone (a gift from Ulrich Technau, Addgene plasmid #67943^83^) with the KikGR fluorophore (from the pCAG::KikGR vector, a gift from Anna-Katerina Hadjantonakis, Addgene plasmid #32608^37^). KikGR reporter expression was driven by the published 2.4kb upstream region of *Elav* from^20^ or the upstream regions of either *SoxC* (2.2kb) or *FoxL2* (2.4kb), with primers in Supplementary File 1. Stable transgenic animals were generated using I-SceI-mediated transgenesis^83^, with the following modified injection mixture: 50ng/μL plasmid concentration, 1X I-SceI buffer (10mM TrisHCl, 10mM MgCl2, 1mM DTT, pH8.8), 0.5mM Patent Blue VF (Sigma, #198218), 0.2U/μL I-SceI (adjusted to 0.4U/μL after 30min at 37°C). All experiments used stably integrated *SoxC>KikGR*, *FoxL2>KikGR* or *Elav>KikGR* 5-8 week-old heterozygote animals (sexually immature adults) raised at 18°C in 12.5ppt artificial seawater in the dark and fed 4 times a week with freshly hatched *Artemia* nauplii.

### Whole animal photoconversion nerve net *in situ* imaging and quantification

Individual *SoxC>KikGR*, *FoxL2>KikGR* or *Elav>KikGR* immature adults were relaxed in 12.5ppt artificial seawater and immobilized by a 1:1 dilution with 70mg/mL MgCl_2_, then transferred between two coverslips, with tape layers to adjust depth to ca. 0.5mm for photoconversion and live confocal imaging. Whole animal KikGR photoconversion was performed with a ZEISS Axio Imager 2 (ZEISS), with a X-Cite 120LED Boost (EXCELITAS) light source at 80% power, with manually tiled regions with 8 sec exposure (emission BP 445/50nm) at 20X magnification. Photoconversion efficiency was confirmed by imaging and flow cytometry (Supplementary Figure 3B) following animal dissociation into single cells. Photoconverted animals were kept in 12.5ppt artificial seawater, with either 1% DMSO or without. For confocal imaging of the body wall nerve net, individual *SoxC>KikGR*, *FoxL2>KikGR* or *Elav>KikGR* adults at 2-, 4-, or 7-days post KikGR photoconversion were immobilized (as above) and imaged at 40X on a Zeiss LSM800 inverted confocal microscope, with 8-10 optical sections per z-stack with 1µm between optical slices with the enhanced green fluorescent protein (eGFP) spectrum used for detecting non-photoconverted KikGR and mCherry spectrum used for the photoconverted KikGR protein, adjusted to remove overlap. For quantification purposes, all images from the same reporter line at 2, 4 or 7 days post-photoconversion were acquired with the same settings. Fig 1B and Fig 3A/C composite extended depth of focus and single slice images and Supplementary Videos 1-9 are for illustration only. All cell segmentations and quantifications were performed on raw z-stack data. For cell quantification, a cellpose 2.0 neural net model^84,85^ was manually trained on randomly sampled individual images slices from *SoxC>KikGR*+, *Elav>KikGR*+ and *FoxL2>KikGR*+ live body wall confocal z-stacks, then used to segment all raw z-stacks at all timepoints post-photoconversion separately on either the red (photoconverted KikGR signal) or green (new KikGR signal) channel for all drivers with the following parameters: diameter: 0, chan2: 0, cellprob_threshold: -0.5,flow_threshold: 0.8, stitch_threshold: 0.25 after optimization. The two sets of cell masks per image (red+ set and green+ set) were then merged into a single set for each z-stack for quantification. Representative videos of the z-stacks with the segmentation are provided in Supplementary Videos 1-9.

### Single cell dissociation and KikGR+ cell sorting

For scRNA-seq sampling, individual *SoxC>KikGR*, *FoxL2>KikGR* or *Elav>KikGR* adults at 2-, 4-, or 7-days post KikGR photoconversion (with initial experiments containing 1% DMSO added to their medium during the timecourse, and not in latter experiments, see Supplementary File 2 and 3) were immobilized with MgCl_2_ as above and sampled below their pharynxes (body without pharynx region or tentacles) to be consistent with our body wall imaging data, then dissociated into single cells according to ^86^, with 1-2 individuals per dissociation. In brief, sample medium was replaced with Calcium Magnesium free seawater (495 mM NaCl, 9.7 mM KCl, 27.6 mM NaHCO_3_, 50 mM Tris–HCl pH 8) with collagenases (0.05mg/mL Liberase TM, Roche), and dissociated by gentle pipetting at room temperature. The cell suspension was filtered (Ø35µm) and stained with live cell dye Calcein Violet AM (ThermoFisher Scientific, 1:2000 dilution of 1µg/µL DMSO stock) or a combination of Calcein Violet and SytoxRed dead cell dye (Thermo Fisher scientific, 1:1000 dilution of a DMSO 5µM stock).

Individual live KikGR+ cells were sorted using a Fluorescence-Assisted Cell Sorter (FACS Aria IIu and III, BD Biosciences) into 384-well plates containing 2µL lysis buffer and barcoded polyT capture oligonucleotides (MARS-Seq protocol^87,88,4^) then frozen at -80°C until processing. FACS gating strategy is shown in Supplementary Fig. 3 and 4; in brief, to sort live KikGR+ cells, cell-sized particles were selected using their FSC-A (logicle-scale) vs SSC-A (logicle-scale) signal (ca. >=3µm in diameter, using sizing beads (SpheroTech) as an absolute size reference). Multiplets were excluded with FSC-W and FSC-H, then Calcein Violet AM-positive live singlets were selected (ex.: 405nm, em.: BP 450/40nm). Initial sorts included a SytoxRed signal: APC channel (ex.: 633nm, em. 660/20nm) and mildly depleted the Calcein Violet^low^/SytoxRed^mid^ population. From live singlets, KikGR+ cells were selected based on their new KikGR signal (FITC-A, ex.:488nm, em. filters: LP 502nm and BP 530nm/30) and photoconverted KikGR (PE-A, ex.: 561nm, em. Filters: BP 582/15nm) signal, with the FITC-A^low^/PE-A^low^ autofluorescent cells excluded, with a negative wild-type (no KikGR) control. A shared KikGR+ gate was used for all *Elav>KikGR*+ and *SoxC>KikGR*+ cells, and for 17/41 of *FoxL2>KikGR+* libraries (Supplementary Fig. 3). For 16/41 *FoxL2>KikGR+* libraries, we depleted its large epithelial cell fraction, as enabled by its specific photoconverted KikGR-positive/low new KikGR signal profile (“Epithelial-depleted KikGR+” sorting). To enrich for green and orange cells, we also sampled only New KikGR^mid/high^ cells for 4 FoxL2>KikGR+ libraries (“New KikGR+” gate). All quantifications of reporter status (green/orange/red cell proportions) from FACS-sorted cells are from cells sorted with the shared KikGR+ gate throughout the paper, with ternary plots (R package ggtern^89^) or barplots (R package ggplot2) visualizations, filtering out clusters with less than 10 sampled cells. For each KikGR+ sorted cell, cell reporter status (i.e. green, orange or red) for all drivers (*Elav*, *FoxL2*, *SoxC*) was assigned using red, orange or green gates defined across all sorted cells (Supplementary Fig. 4). Cell identifiers were corrected for two sorted 384-wells plates (4 libraries: RP0885_E/O, RP0886_E/O), as the flow cytometer mechanical arm moved by one column prior to the sort. For each library sorted with the KikGR+ gate and used for color reporter analysis, matched flow cytometry data was validated through its correlation between FSC-A signal (a cell volume proxy) and total UMI per cell.

### Single cell RNA-Seq analysis

We generated 84 scRNA-Seq libraries according to published MARS-Seq protocols^4,87,88^. In brief, all plates were processed with the same protocol, mRNA was reverse transcribed into cDNA with a barcoded oligodT capture oligonucleotide in the lysis buffer containing the cell barcode and unique molecular identifier. Exonuclease treatment (ExoI, NEB) was used to remove unused oligodT nucleotides. Pools of cDNA were then amplified by *in vitro* transcription (T7 polymerase), then fragmented. Each RNA fragment was ligated to an oligonucleotide containing a library pooling barcode with a T4 ssDNA:RNA ligase before reverse transcription. Final libraries were obtained from PCR amplification with 17 cycles. Library fragment distributions and concentrations were assessed by Tapestation 4150 (Agilent) and Qubit3.0 (Invitrogen). scRNA-Sseq libraries were pooled equimolarly and sequenced in paired-end mode on a NextSeq550 (Illumina) using a high output 75 cycles kit (up to 32 libraries per run with 190 cells per half-plate library). Median sequencing depth per library was 13.4M reads (for 190 sorted cells). Fastq files were reformatted to the following read structure for compatibility with STARSolo: Read1: mRNA fragment; Read2: 4 nucleotides (nt) library barcode, 7 nt well barcode (11nt total for the cell barcode), 8 nt UMI barcode. Reads were mapped and demultiplexed using STARSolo, with the following parameters --soloType CB_UMI_Simple --soloUMIlen 8 --soloCBlen 11 --soloUMIstart 12 --soloCBstart 1 --soloCBmatchWLtype 1MM --soloUMIdedup Exact; with the cell barcode list for each sequencing run as the soloCBwhitelist. Paired-end fastq files per half-plate library are provided in the GEO archive (GSE288441), along with cell barcode lists for each pool of demultiplexing-compatible libraries.

Reads were mapped to the jaNemVect1.1 *Nematostella vectensis* genome assembly (from the Darwin Tree of Life (DToL) project – Wellcome Open Research^90^), with the KikGR sequence added^37^, using the genes models from the RefSeq *Nematostella vectensis Annotation Release 101* from NCBI (GCF_932526225.1, NCBI) (in GEO archive GSE288441). Of the 84 libraries, a total of 19, 24, and 41 correspond to the *Elav*, *SoxC*, and *FoxL2* drivers, respectively, with an average of 2131 median RNA molecules recovered per library. Cells were filtered based on RNA molecules recovery (excluding cells with ≤300 or ≥13000 recovered RNA molecules), and low quality clusters (low UMIs, no specific marker gene, large fraction of stress-related genes and ribosomal proteins) were filtered out, as well as clusters with cells from one biological dissociation, only from DMSO+ samples or from a single sequencing run. After filtering, 12352 cells were retained, comprising 78% of sorted cells. Filtered and unfiltered UMI tables are available in the GEO archive (GSE288441). Libraries description with biological samples, experimental condition and gene recovery statistics are in Supplementary File 2. Library RNA molecule recovery is shown in Supplementary Fig. 9. We then used Seurat v5 with SCTransform v2 normalization for clustering, using the top 4000 variable genes (excluding the KikGR transgene)^91–93^, with the top 65 PCA components used and a 2.4 clustering resolution. Clusters were annotated based on literature-curated markers (from ^5,13,15,17,18,34,42,44,45,48,94^). The *Elav* subclustering (Fig. 2) was processed similarly, with only *Elav>KikGR*+ cells included.

Due to the multiple genome assemblies and gene models used in the literature, gene annotations were transferred to the GCF_932526225.1 gene models (*jaNemVect1.1* genome assembly, gene identifiers starting with LOC) using either BLASTP of protein sequences, primerBLAST of *in situ* hybridization probes from the original publications or best reciprocal BLASTP hits; as listed in Supplementary File 1, with the GCF_932526225.1 gene annotation used by default for all other genes. Transcription factors were predicted with AnimalTFDB^95^, with bZIP transcription factors named from their families from^94^, and bHLH transcription factors annotated from^11,55^. Cell cycle module genes (22 G2/M and 29 S-phase genes) were identified in *Nematostella* by mapping Seurat’s mouse cell cycle markers using eggNOG mapper annotations. The distribution of cell identities across libraries was consistent across biological samples for each driver and fluorescence-activated cell sorting (FACS) gating strategy (Supplementary Fig. 9). Gene expression across clusters is visualized using DotPlot from SCTransform data, with expression values normalized to the maximum expression observed across clusters, and dot size representing the percentage of cells expressing the gene in each cluster.

To investigate differential neural effector gene usage between peptidergic neuron clusters (18 clusters) and late cnidocytes (1 cluster), we annotated genes coding for Class A GPCRs, mGluR-like receptors, and various ion channels based on PFAM domains using eggNOG mapper v2. The PFAM domains used included 7tm_1 and 7tm_3 for GPCRs (respectively Class A, and mGluR-like), as well as ASC, Anoctamin, NT_gated_channel, iGluR, PKD_channel, Lig_chan-Glu_bd, and Neur_chan_LBD for ion channels. De-orphanized neuropeptide GPCR receptors were further annotated from ^15^. To assess the differential usage of neural-specific genes, we excluded lowly expressed genes (defined as those with fewer than 50 UMI across neural clusters) and genes whose highest expression occurred in non-neural clusters. Given the presence of multiple peptidergic clusters but only a single late cnidocyte cluster, we systematically compared the late cnidocyte cluster to the peptidergic cluster with the highest expression of each effector gene. The expression matrix was aggregated using Seurat’s AggregateExpression function, with a scale factor of 1000. We calculated the log2 fold change in expression between the late cnidocytes and the peptidergic cluster with maximal effector expression, plotting this against the average expression between the two clusters for 499 genes across six categories (Source Data File 6).

For the cnidocytes developmental trajectory inference, cnidocytes (clusters 6 and 14) were subsetted from the global scRNA-seq clustering and processed in Seurat with standard normalization to generate a UMAP embedding. Pseudotime analysis was performed using slingshot^96^ on the UMAP coordinates. Gene expression along pseudotime was then estimated using a generalized additive model with the gam function from the mgcv R package.

Weighted Gene Co-expression Network Analysis (as implemented in the WGCNA R package) was used to identify 48 transcriptional modules in the global clustering^54^. Gene expression data from the Seurat object were filtered to retain genes with at least 50 UMIs, and aggregated expression was computed per cluster with log normalization and a scale factor of 1000. A weighted gene co-expression network was then constructed using a soft-threshold power of 6. The adjacency matrix was transformed into a topological overlap matrix (TOM), and hierarchical clustering was performed. Modules were identified via dynamic tree cutting (deepSplit = 3, minClusterSize = 20), then merged based on eigengene correlations, using a dissimilarity threshold of 0.25. Module eigengenes were computed and gene-module membership was determined by correlating gene expression with module eigengenes across clusters. A low (0.35) and high (0.5) threshold were used to visualize soft module membership for transcription factors. Gene module expression across clusters (Supplementary Figure 15) was computed using Seurat’s AddModuleScore function with each module composed of genes with module membership ≥0.5 (stringent threshold), and each gene only associated to its top scoring module (Source Data File 6). These thresholded gene modules were then used to visualize the network of module top co-expression relationships, derived from the average pairwise TOM values, by filtering out module-module average scores below 0.002.

### Reanalysis of RNA-Seq data

Upregulated and downregulated genes from the *PouIV-KO* and *Prdm14d>GFP*+ RNA-Seq datasets, as reported in the original publications^13,18^, were converted to GCF_932526225.1 gene identifiers using best reciprocal BLASTp hits (Supplementary File 1). Gene set enrichment analysis was then performed using the GSEA R package, with cluster gene markers from the FindAllMarkers in Seurat function on the global clustering (with an adjusted p-value threshold of 0.01) used for the ’pathways’ lists (26-1265 genes per cluster-associated set) (Source Data File 7).

## Acknowledgments

We thank Sébastien Bastide for data analysis discussions and comments on the manuscript. We thank Caroline Kaiser, Ilaria Rebay, Paschalis Kratsios and Urs Schmidt-Ott, and members of the Marlow lab for discussions and comments on the manuscript. We thank Colette Plessier for the *Nematostella* silhouettes. We thank Baptiste Saudemont for MARS-Seq training, Sophie Novault and Sandrine Schmutz for FACS training and expert advice. We thank the Flow Cytometry Core Facility in Institut Pasteur (Paris, France) for BD FACS AriaIII access. We thank David Leclerc, Mike Olson and Bert Ladd of the University of Chicago Cytometry and Antibody Technology Facility for BD FACS AriaII training and advice *(Facility RRID: SCR_017760*). We thank Marie-Pierre Mailhé for assistance with animal care. We thank the Developmental and Stem Cell Biology Department of Institut Pasteur (Paris, France) for Zeiss LSM800 confocal microscope access. We thank Matt Kaufman for image processing advice. We thank the University of Chicago’s Research Computing Center for HPC access for image processing. We thank the Ecole Normale Supérieure de Lyon and the Ministère de l’Enseignement Supérieur et de la Recherche (France) for funding (to F.P.).

## Author Contributions

H.M. conceived the study. H.M. and F.P. designed experiments and interpreted the data. F.P. performed experiments, developed and implemented data analyses, generated figures, and wrote the initial draft. H.M. and F.P. wrote and edited the manuscript. H.M. supervised the project and acquired funding.

