## Supplementary Figures and Legends for "Cellular and transcriptional trajectories of neural fate specification in sea anemone uncover two modes of adult neurogenesis"

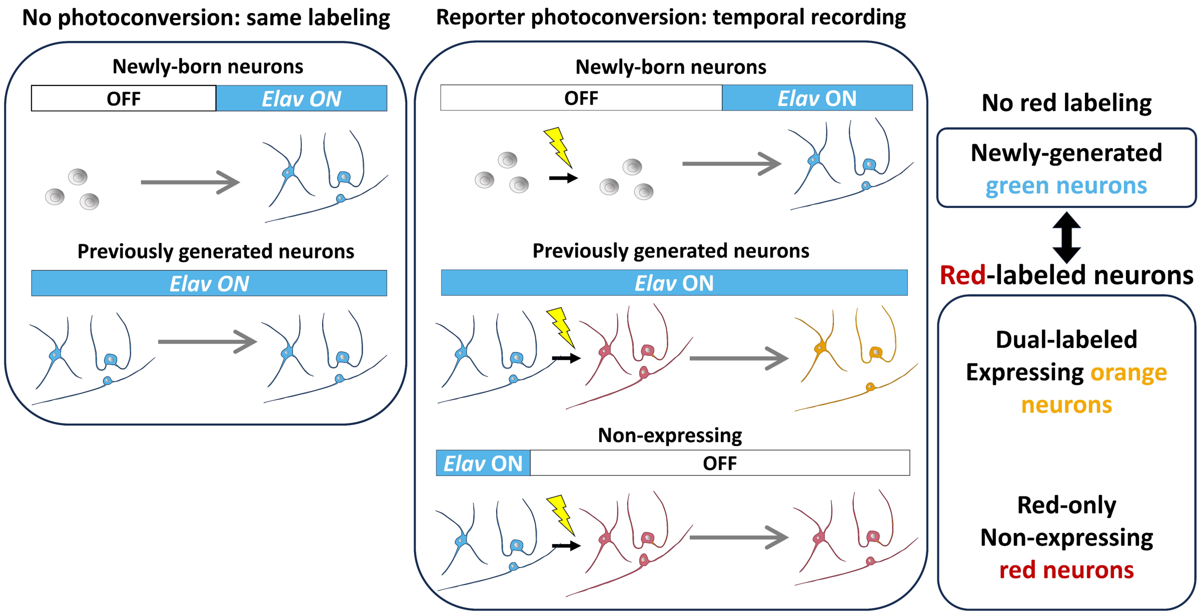


**Supplementary Figure 1: Discriminating adult-born neurons by photoconversion-based temporal recording of reporter expression**

Reporter driver with no photoconversion (**left**) does not discriminate between previously expressing cells and newly expressing cells. A photoconvertible driver strategy (**right**) can label all previously expressing neurons at time 0 and then be used to track newly-expressing-only neurons by their presence of the new reporter protein (green signal, in blue here) and the lack of the photoconverted red labeling. Within the red-labeled neurons, red non-expressing progeny cells of previously expressing cells can be further distinguished from expressing progeny cells (contain both the photoconverted red label and the new green label, represented as orange cells throughout).

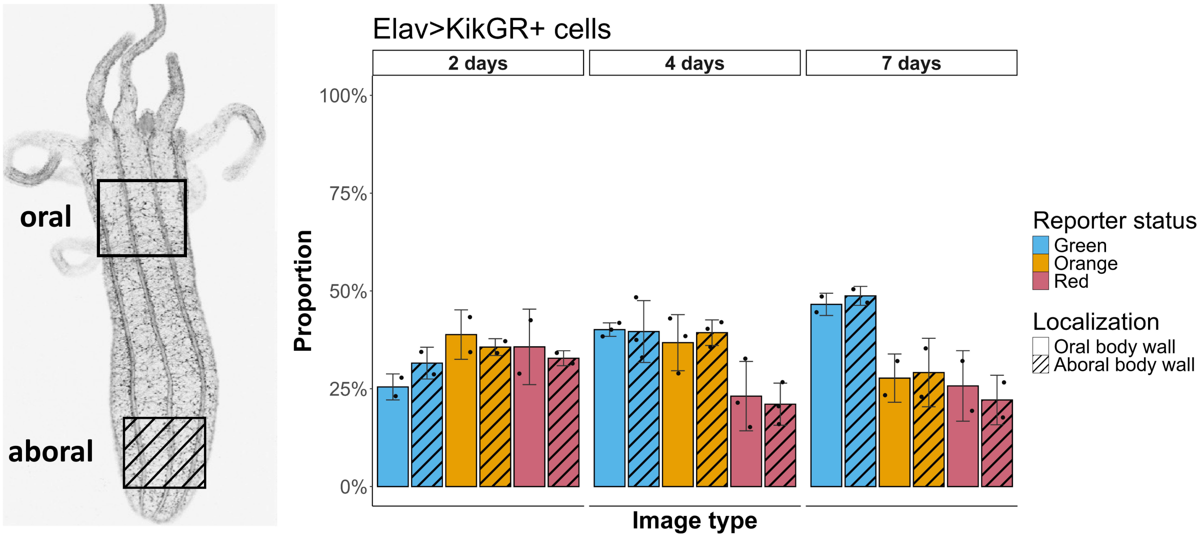


**Supplementary Figure 2: Spatio-temporal dynamics of neurogenesis across the primary body axis**

**A** Quantification of the proportion of *Elav>KikGR+* detected cells in the nerve net by reporter status, time post photoconversion and primary body axis localization. Cell proportion is normalized to all cells detected within each confocal z-stack image, with the red, orange and green (represented in blue) cell fraction. The relative fraction of green neurons (newly-expressing only neurons arising from non-expressing cells) increases with the time post-photoconversion across both the oral (no hachures) and aboral (with hachures) locations. n=2-3 animals per condition, with dots for the individual image quantifications (mean ±S.D.). See Supplementary Videos 1-2.

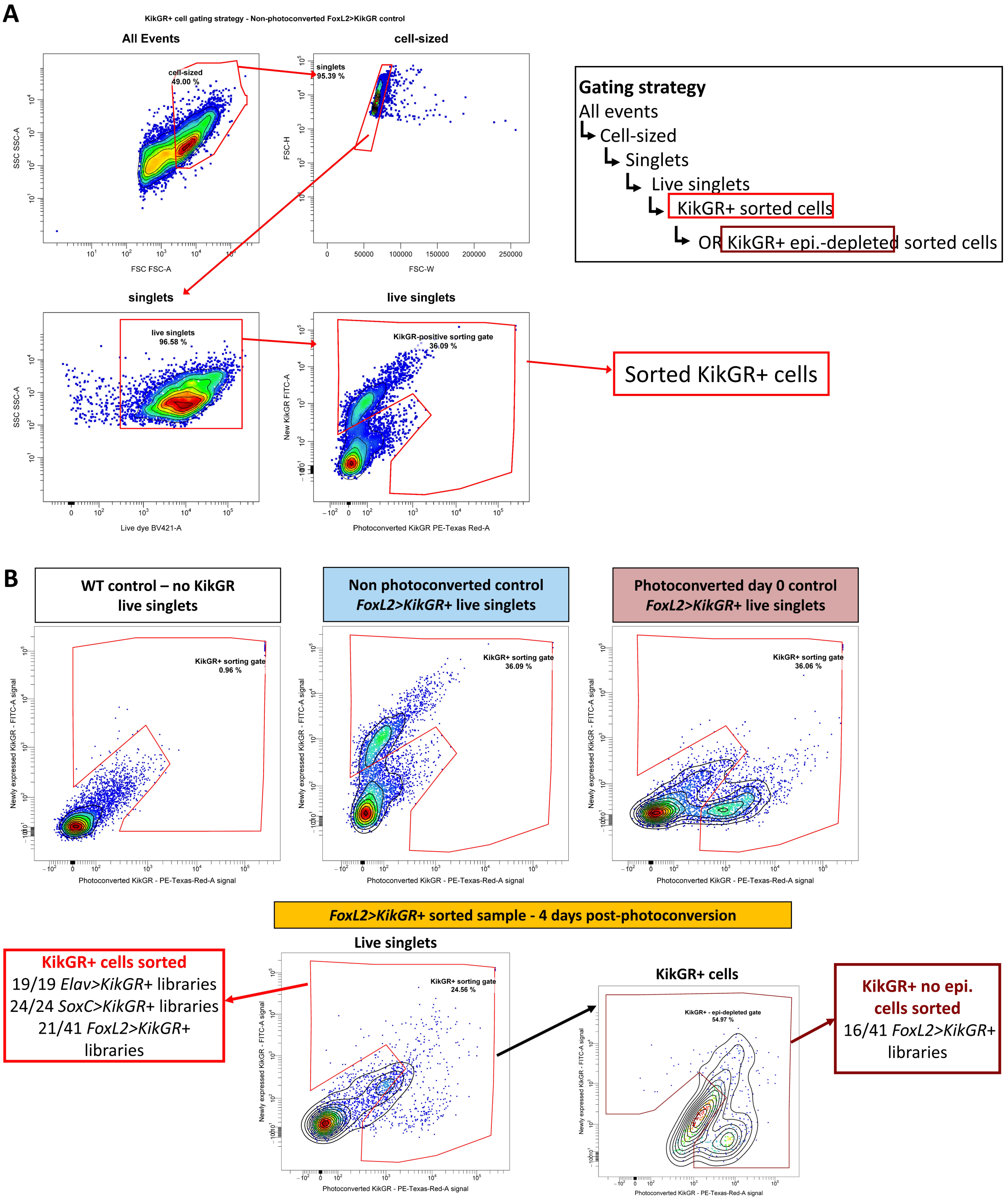


**Supplementary Figure 3: KikGR+ cell flow cytometer gating strategy**

**A** Fluorescence-Activated Cell Sorter (FACS) strategy used to sample *KikGR*+ cells from adult body (excluding pharyngeal and tentacles regions) cell suspension for *Elav*, *FoxL2* and *SoxC* drivers. From all events, cells sized particles (approximately ≥3µm in diameter, from sizing beads) are selected, then multiplets are excluded based on their FSC-W/FSC-W signal and Calcein-positive cells are retained (live singlets fraction). *KikGR*+ cells are selected, with autofluorecent cells excluded (see **B**). **B** From live singlets, the KikGR+ sorting gate is defined with WT (wild-type, no KikGR reporter), non-photoconverted (all 'green' cells) and fully-photoconverted (day 0, all 'red' cells) control samples. Autofluorescent cells (low PE-Texas-Red-A/low FITC-A double-positive signal) are excluded from the KikGR+ gate using the WT control sample. The same KikGR+ sorting gate is used for all *Elav>KikGR*+ and *SoxC>KikGR*+ cells at all timepoints post-photoconversion. *FoxL2>KikGR*+ cells are sorted with either this *KikGR*+ gate or with the '*KikGR*+ no epi.' gate to deplete the epidermis epithelial cell fraction and enrich for populations of interest (progenitor and neurons). All quantification of reporter color status for all drivers throughout the study (i.e. green/orange/red cell fractions) only use cells sorted with the *KikGR*+ sorting gate.

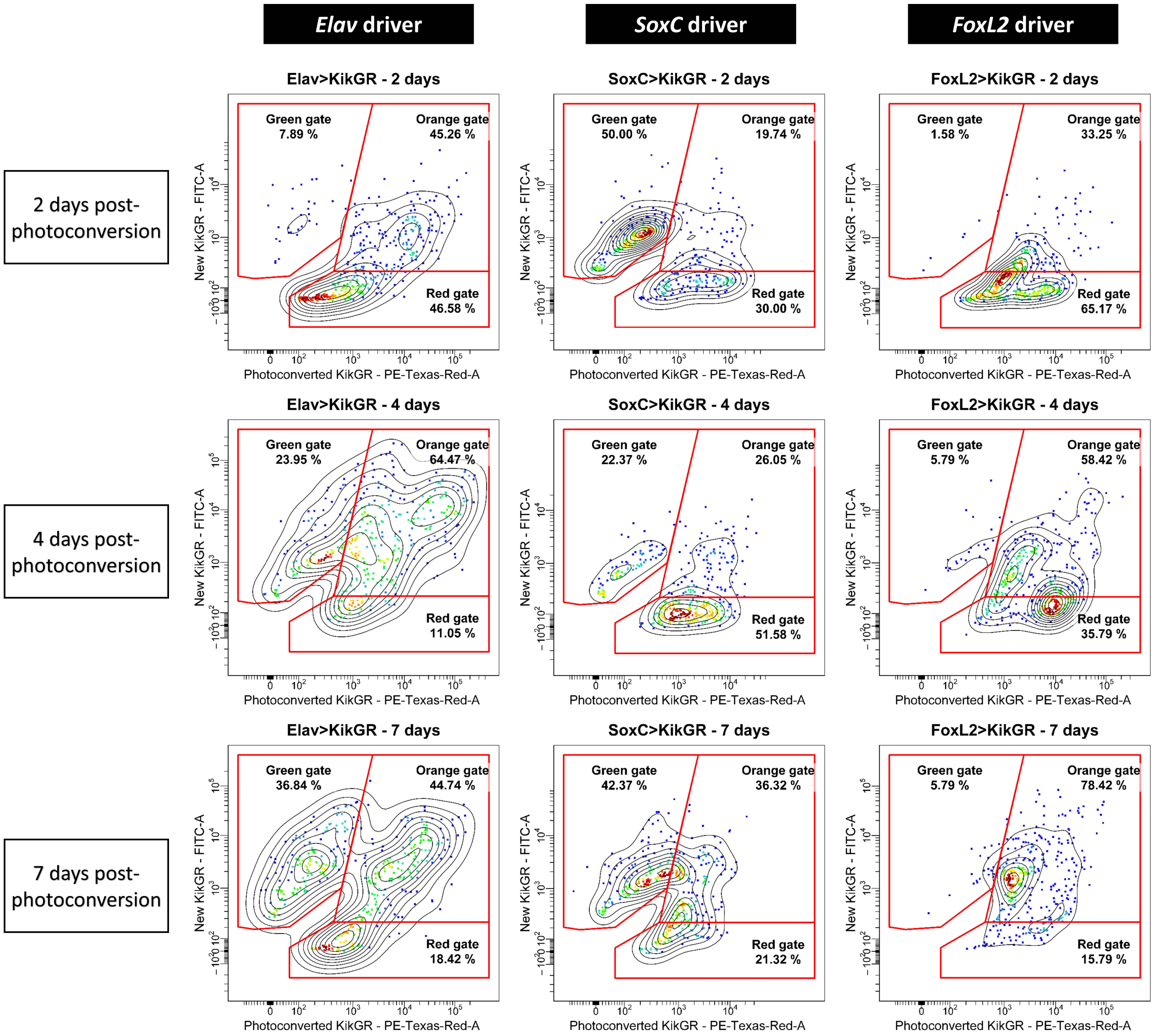


**Supplementary Figure 4: Reporter status attribution ('green', 'orange' or 'red' cell)**

"Green", "orange" and "red" gates are defined for all KikGR+ sampled cells (KikGR+ sorting gate, see Supplementary Fig. 3) at 2,4, or 7 days post-photoconversion for the *Elav*, *SoxC* and *FoxL2* drivers. Representative data from 380 sorted cells (2 half-plate libraries) shown for each driver and timepoint post-photoconversion.

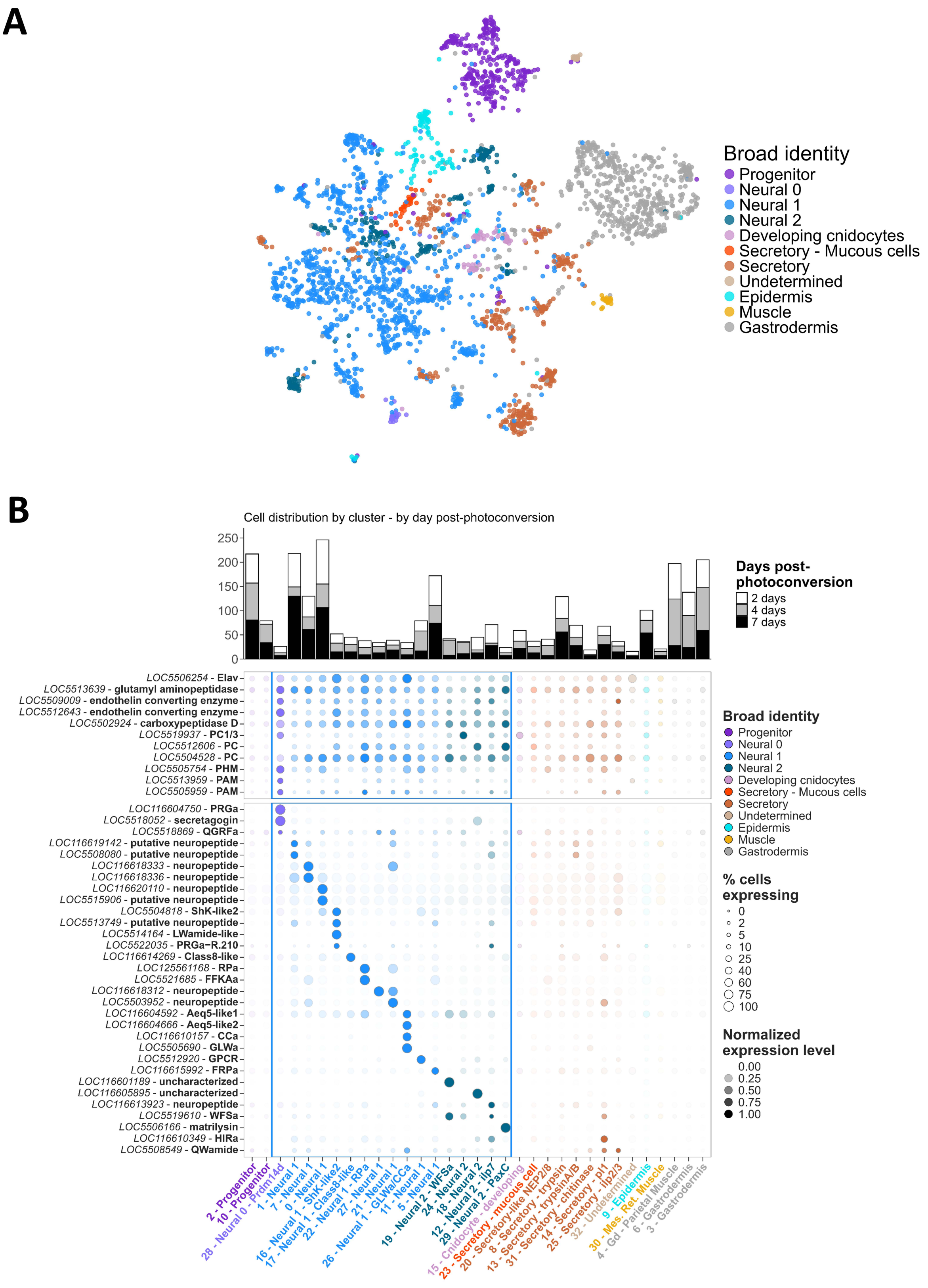

A
Broad identity
Progenitor
Neural 0
Neural 1
Neural 2
Developing cnidocytes
Secretory - Mucous cells
Secretory
Undetermined
Epidermis
Muscle
Gastrodermis
B
Cell distribution by cluster - by day post-photoconversion
250
200
150
100
50
0
Days post-photoconversion
2 days
4 days
7 days
LOC5506254 - Elav
LOC5513639 - glutamyl aminopeptidase
LOC5509009 - endothelin converting enzyme
LOC5512643 - endothelin converting enzyme
LOC5502924 - carboxypeptidase D
LOC5519937 - PC1/3
LOC5512606 - PC
LOC5504528 - PC
LOC5505754 - PHM
LOC5513959 - PAM
LOC5505959 - PAM
LOC116604750 - PRGa
LOC5518052 - secretagogin
LOC5518869 - QGRFa
LOC116619142 - putative neuropeptide
LOC5508080 - putative neuropeptide
LOC116618333 - neuropeptide
LOC116618336 - neuropeptide
LOC116620110 - neuropeptide
LOC5515906 - putative neuropeptide
LOC5504818 - ShK-like2
LOC5513749 - putative neuropeptide
LOC5514164 - LWamide-like
LOC5522035 - PRGa−R.210
LOC116614269 - Class8-like
LOC125561168 - RPa
LOC5521685 - FFKAa
LOC116618312 - neuropeptide
LOC5503952 - neuropeptide
LOC116604592 - Aeq5-like1
LOC116604666 - Aeq5-like2
LOC116610157 - CCa
LOC5505690 - GLWa
LOC5512920 - GPCR
LOC116615992 - FRPa
LOC116601189 - uncharacterized
LOC116605895 - uncharacterized
LOC116613923 - neuropeptide
LOC5519610 - WFSa
LOC5506166 - matrilysin
LOC116610349 - HIRa
LOC5508549 - QWamide
Broad identity
Progenitor
Neural 0
Neural 1
Neural 2
Developing cnidocytes
Secretory - Mucous cells
Secretory
Undetermined
Epidermis
Muscle
Gastrodermis
% cells expressing
0
2
5
10
25
40
60
75
100
Normalized expression level
0.00
0.25
0.50
0.75
1.00
2 - Progenitor
10 - Progenitor
28 - Neural 0 - Prdm14d
1 - Neural 1
7 - Neural 1
0 - Neural 1
16 - Neural 1 - ShK-like2
17 - Neural 1 - Class8-like
22 - Neural 1 - RPa
27 - Neural 1
26 - Neural 1 - GLWa/CCa
21 - Neural 1
11 - Neural 1
5 - Neural 1
19 - Neural 2 - WFSa
24 - Neural 2
12 - Neural 2
18 - Neural 2 - Ilp7
29 - Neural 2 - PaxC
15 - Cnidocyte - developing
23 - Secretory - mucous cell
20 - Secretory-like NEP2/8
8 - Secretory - trypsin
13 - Secretory - trypsinA/B
31 - Secretory - chitinase
14 - Secretory - pl1
25 - Secretory - ilp2/3
32 - Undetermined
30 - Mes. Ret. Muscle
9 - Epidermis
4 - Gd - Parietal Muscle
6 - Gastrodermis
3 - Gastrodermis

**Supplementary Figure 5: Neuropeptide processing genes and neuropeptide precursors expression across all *Elav>KikGR*+ sorted cells**

**A** UMAP projection of *Elav>KikGR*+ cells, colored by cell class identity. **B** Distribution of 2,4,7 days post-photoconversion *Elav>KikGR*+ cells across all clusters (**top**) and marker gene expression for peptidergic processing enzymes and neuropeptide precursors across all *Elav>KikGR*+ cells (same clustering as in A and Fig. 2C) (**bottom**). Cells from different timepoint post-photoconversion are broadly distributed across molecular identities, which consistent with steady-state sampling during growth. See Supplementary Fig. 9 for the cell identity distribution for all scRNA-seq libraries. Marker genes are the same as in Fig. 2C with expression normalized to maximum gene expression across all clusters. Dot size represents the percentage of cells expressing the gene in each cluster. Neural clusters shown in Fig. 2C are highlighted with the blue rectangle. Data in Source Data File 2.

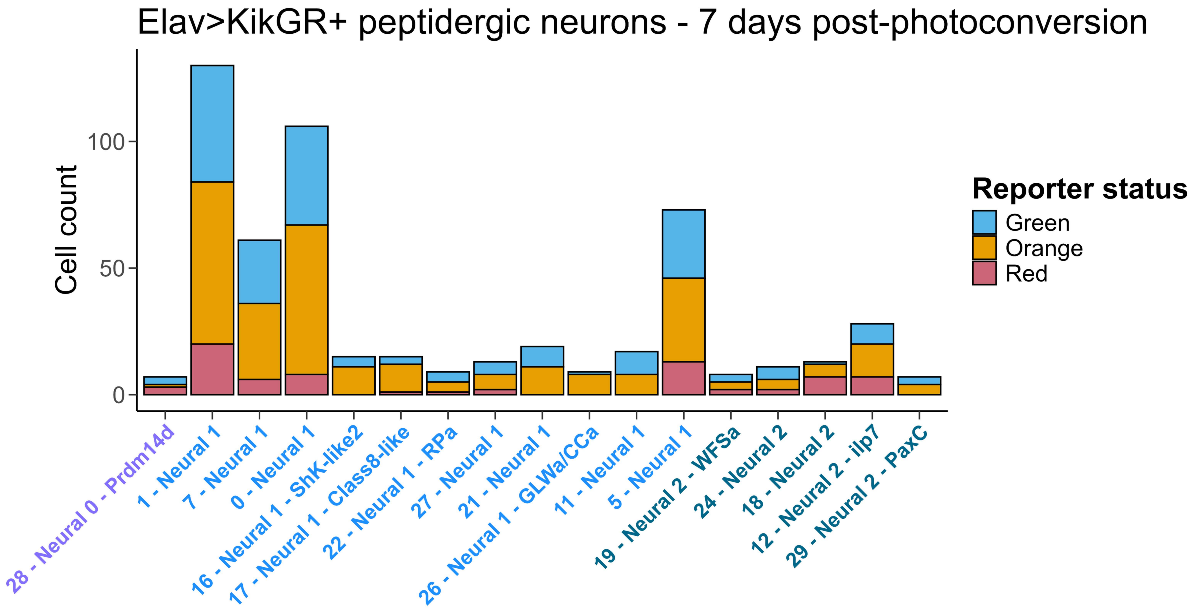


**Supplementary Figure 6: Reporter distribution in Elav>KikGR+ peptidergic neurons after one week**

The smaller fraction of red neurons (13%, 76/569 cells) does not appear to be exclusive to specific peptidergic subtypes, although sampling limits our analysis for the lower-abundance subtypes.

**A**

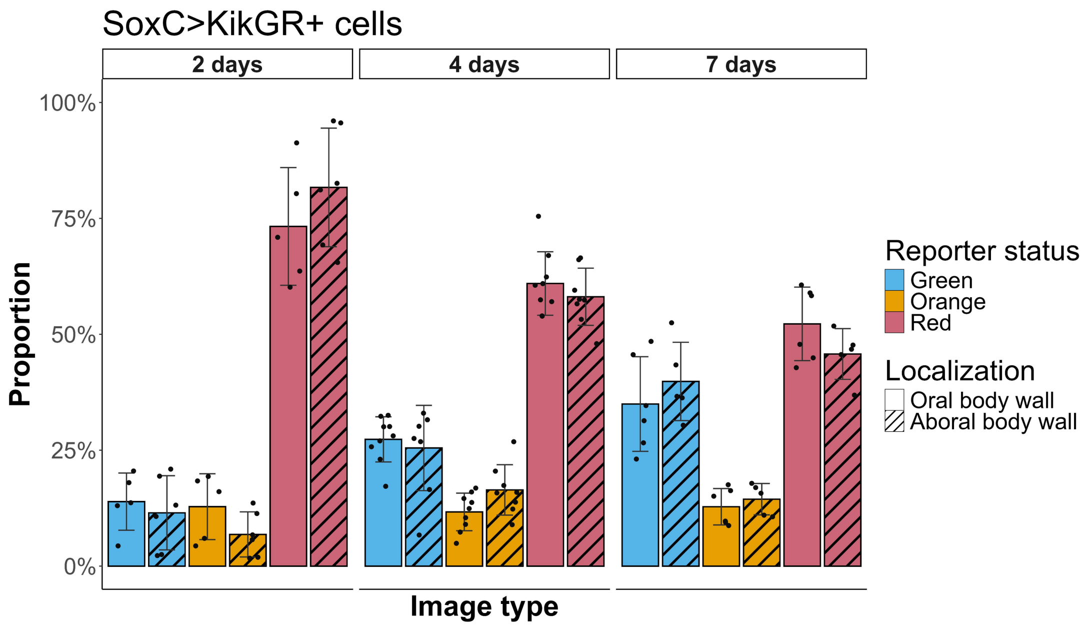


**B**

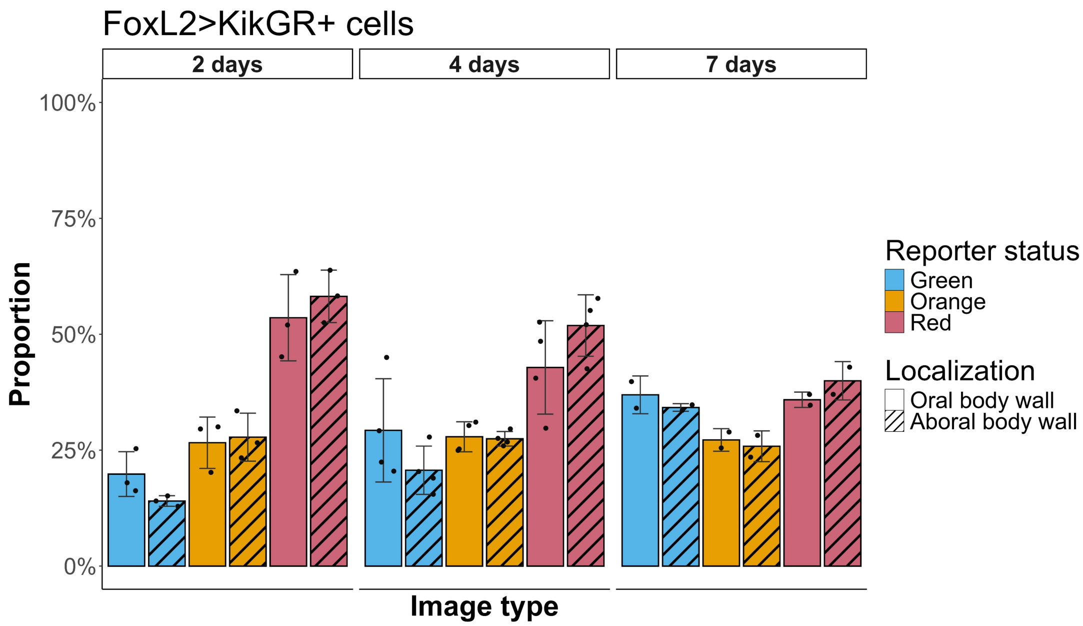


**Supplementary Figure 7: *SoxC* and *FoxL2* drivers oral and aboral body wall reporter dynamics cellular quantification**

**A-B** Proportions of detected *SoxC>KikGR+* (**A**) and *FoxL2>KikGR+* (**B**) cells in the nerve net region by reporter status, time post reporter photoconversion and localization. Cell proportion is normalized to all cells detected within each z-stack image, with the red, orange and green (in blue) cell fractions. The fraction of green cells increases with the days post-photoconversion across both the oral (no pattern) and aboral (with hachures) areas. n=2-7 animals per reporter and post-photoconversion timepoint, with dots for individual image quantifications (mean ±S.D.). See Supplementary Videos 3-9.

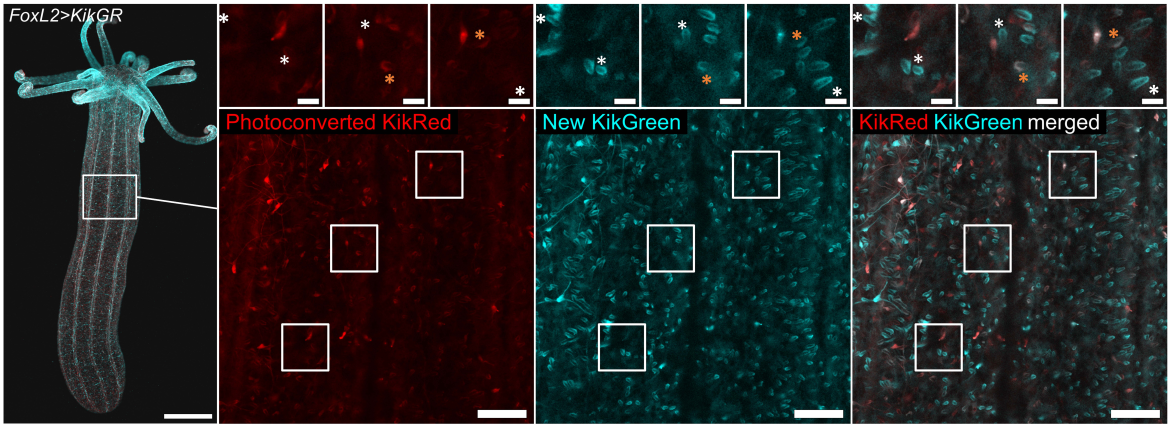


**Supplementary Figure 8: Epidermis-embedded mature cnidocytes, epithelial cells and putative sensory neurons are labeled by the *FoxL2* driver**

*FoxL2>KikGR*+ oral body wall at 7 days post-photoconversion, epidermis plane of focus from the same image as Fig 3C. 5X (**left**) and 40X (B) view with photoconverted KikGR (in red), new KikGR signal (in blue), and merged (**right**). Green, and orange mature cnidocytes (white, orange and magenta asterisks, respectively) are interspersed with orange epithelial epidermis cells and red and orange epithelial sensory neurons. Scale bars are 500 µm (left), 50 µm (right), and 10 µm in the insets. See Supplementary Video 8.

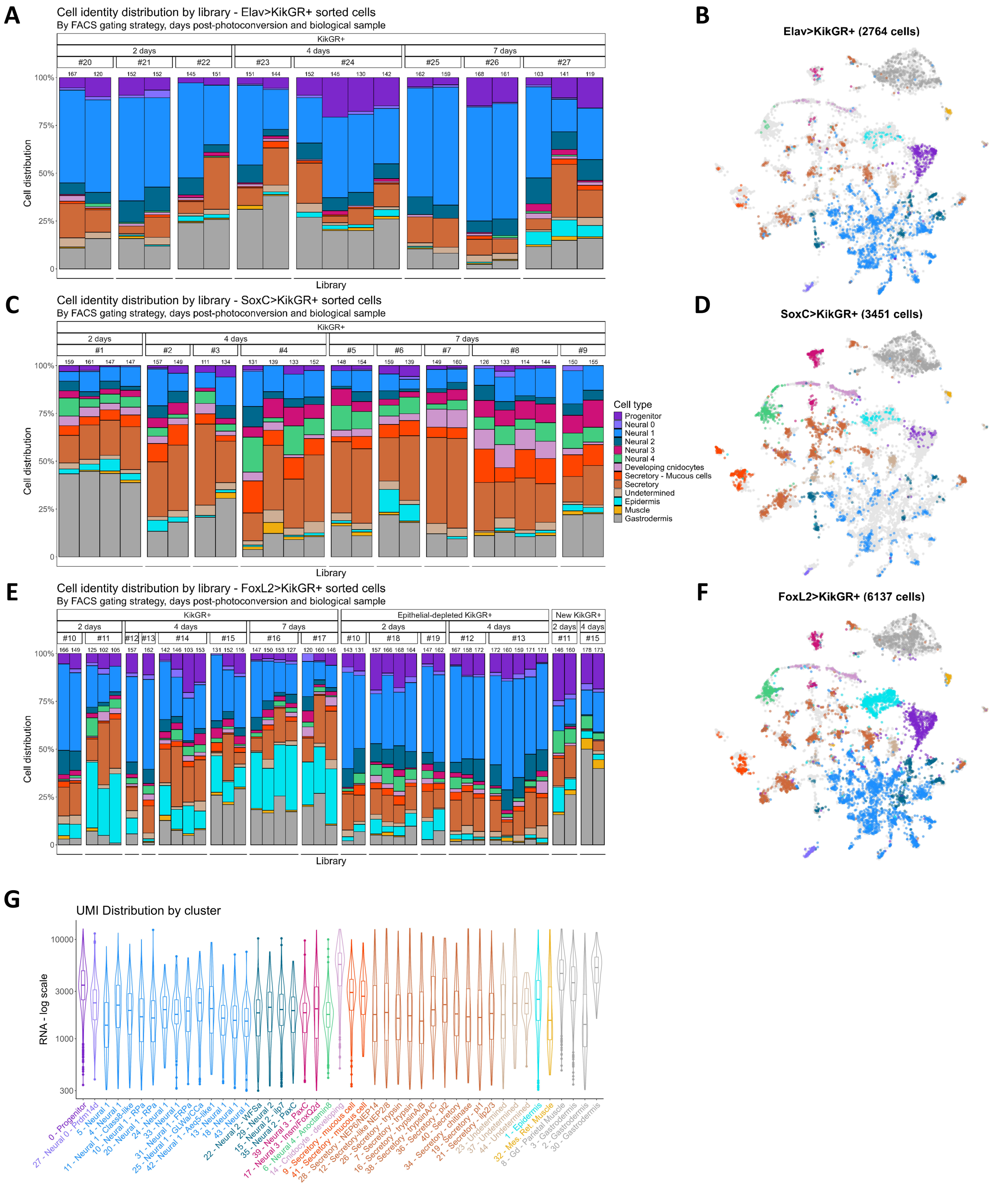


### Supplementary Figure 9: scRNA-Seq QC: Cell identity distribution by scRNA-Seq library

**A,C,E** Cell identity distribution by scRNA-Seq library by FACS sorting strategy, day post reporter photoconversion and biological sample and driver (**A**: *Elav* driver, **C** *SoxC* driver, **E** *FoxL2* driver). We find similar distribution of broad cell identity classes within each driver line, from the distinct biological cell suspensions recovered. The number of cells recovered after filtering from each library is indicated above each barplot. **B,D,F** UMAP with all KikGR+ cells from the global clustering, colored by cells recovered from the *Elav* (**B**), *SoxC* (**D**) or *FoxL2* (**F**) driver. **G** Recovered RNA molecules across all clusters. Broad cell identity colors as in A-F.

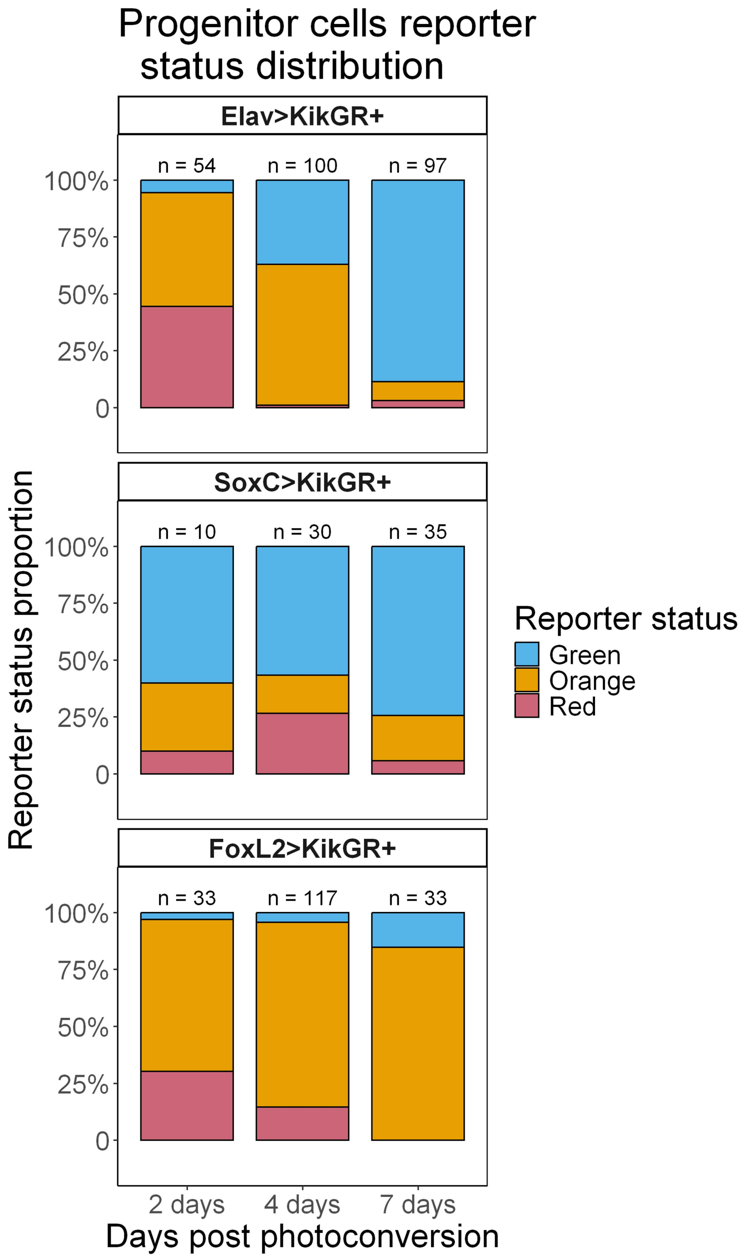


**Supplementary Figure 10: Progenitor reporter dynamics from the *Elav*, *SoxC* and *FoxL2* drivers highlighting exit-bound (*Elav*+ or *SoxC*+) cells within this compartment.**

Same data as represented by ternary plots in Fig. 4B, with the number of cells recovered for each driver and recording timepoint post-photoconversion indicated above. Green cells are represented in blue.

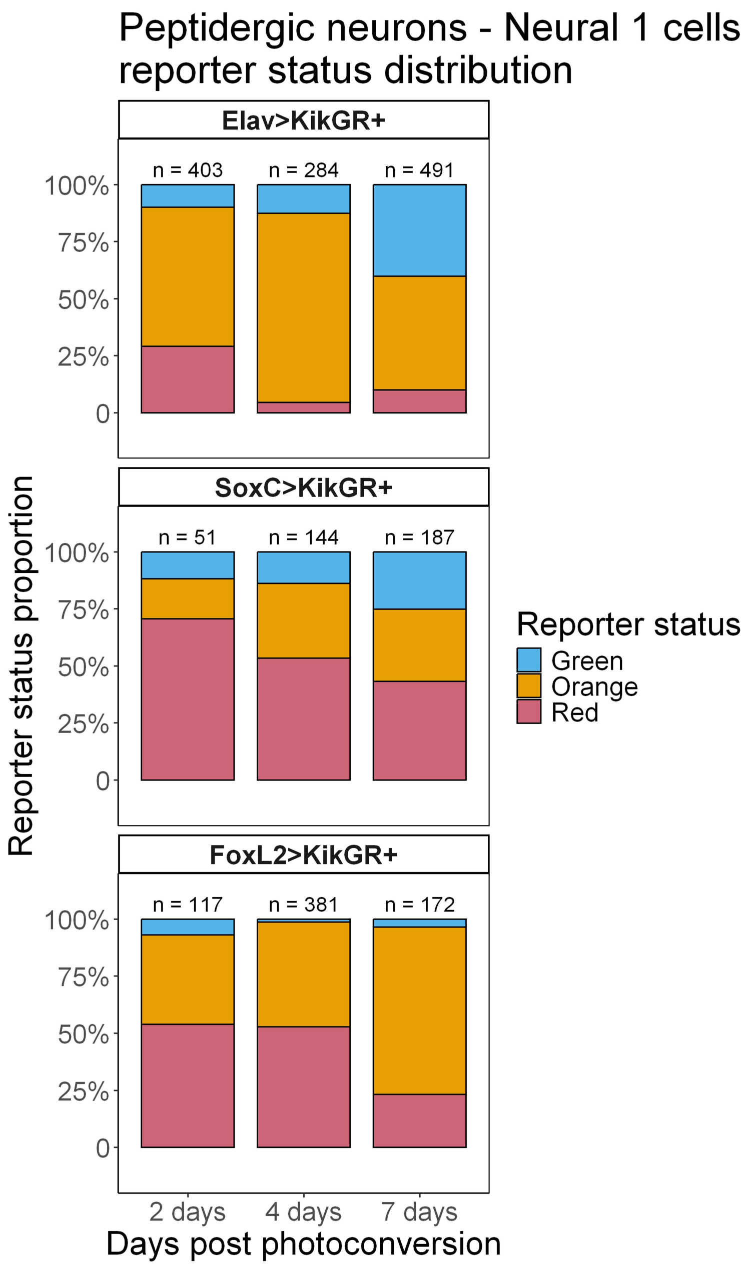


**Supplementary Figure 11: Peptidergic neurons (Neural 1 class) reporter dynamics from the *Elav*, *SoxC* and *FoxL2* drivers**

Same data as represented by ternary plots in Fig. 5B, with the number of cells recovered for each driver and recording timepoint post-photoconversion indicated above. Green cells are represented in blue. The large fraction of red peptidergic neurons progeny cells from the *SoxC* and *FoxL2* drivers is consistent with these arising from the multipotent progenitor pool.

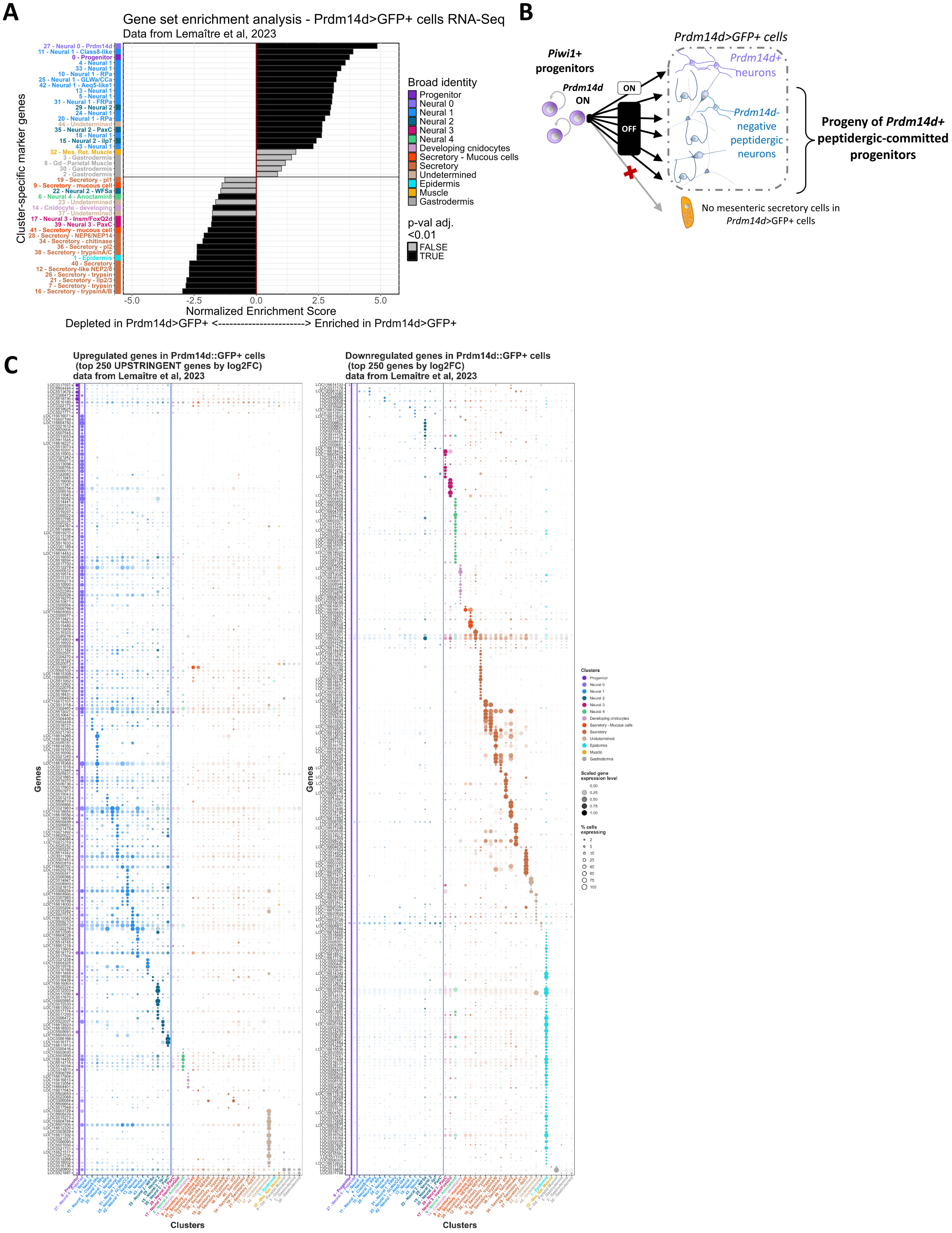
A
Gene set enrichment analysis - Prdm14d>GFP+ cells RNA-Seq
Data from Lemaître et al, 2023
Cluster-specific marker genes
Broad identity
Progenitor
Neural 0
Neural 1
Neural 2
Neural 3
Neural 4
Developing cnidocytes
Secretory - Mucous cells
Secretory
Undetermined
Epidermis
Muscle
Gastrodermis
p-val adj. <0.01
FALSE
TRUE
-5.0
-2.5
0.0
2.5
5.0
Normalized Enrichment Score
Depleted in Prdm14d>GFP+ <-----------------------> Enriched in Prdm14d>GFP+
B
Piwi1+ progenitors
Prdm14d ON
ON
OFF
Prdm14d>GFP+ cells
Prdm14d+ neurons
Prdm14d- negative peptidergic neurons
Progeny of Prdm14d+ peptidergic-committed progenitors
No mesenteric secretory cells in Prdm14d>GFP+ cells
C
Upregulated genes in Prdm14d::GFP+ cells (top 250 UPSTRINGENT genes by log2FC) data from Lemaître et al, 2023
Downregulated genes in Prdm14d::GFP+ cells (top 250 genes by log2FC) data from Lemaître et al, 2023
Genes
Clusters
Scaled gene expression level
% cells expressing

**Supplementary Figure 12: Reanalysis of *Prdm14d* driver RNA-Seq data supports a peptidergic-committed progenitor lineage**

**A** Gene set enrichment analysis (GSEA) assessing the enrichment of cluster marker genes in the *Prdm14d>GFP*+ RNA-Seq dataset. Genes enriched in *Prdm14d>GFP*+ cells (Normalized Enrichment Score >0) overlap with marker genes expressed in peptidergic clusters (Neural1 and 2 class, which do not express *Prdm14d*), while genes depleted in *Prdm14d>GFP*+ cells (Normalized Enrichment Score <0) overlap with other differentiated cell type markers, including those of secretory cells, and cnidocytes. **B** Model of *Prdm14d*+ progenitors giving rise to peptidergic neurons but not to secretory cells, as inferred from the reanalysis of the *Prdm14d>GFP*+ RNA-Seq dataset. *Prdm14d* is endogenously expressed in *Piwi1*+ progenitors, and in one peptidergic neuron population (Neural 0, *Prdm14d*+ neurons), but not in other peptidergic neurons (Neural 1 and 2) (Fig. 3F). Neural 1 peptidergic neuron markers are expressed in *Prdm14d>GFP*+ cells (in **A** and **C**). **C** Expression of the top 250 upregulated genes (stringent filtering from the original dataset, **left**) and top 250 downregulated genes (**right**) from *Prdm14d>GFP*+ cells across clusters (same clustering as in Fig. 3). Genes upregulated in GFP+ cells from the *Prdm14d* driver are expressed in progenitors (which express *Prdm14d*) and across both *Prdm14d*+ peptidergic neurons (Neural 0) and *Prdm14d*-negative peptidergic neurons (Neural 1 and 2). Genes downregulated in the *GFP*+ cells are specific to other differentiated cell types, including cnidocytes, secretory cells, epidermis epithelial cells, which suggest that are not part of the shared peptidergic-committed *Prdm14d*+ progenitor lineage.

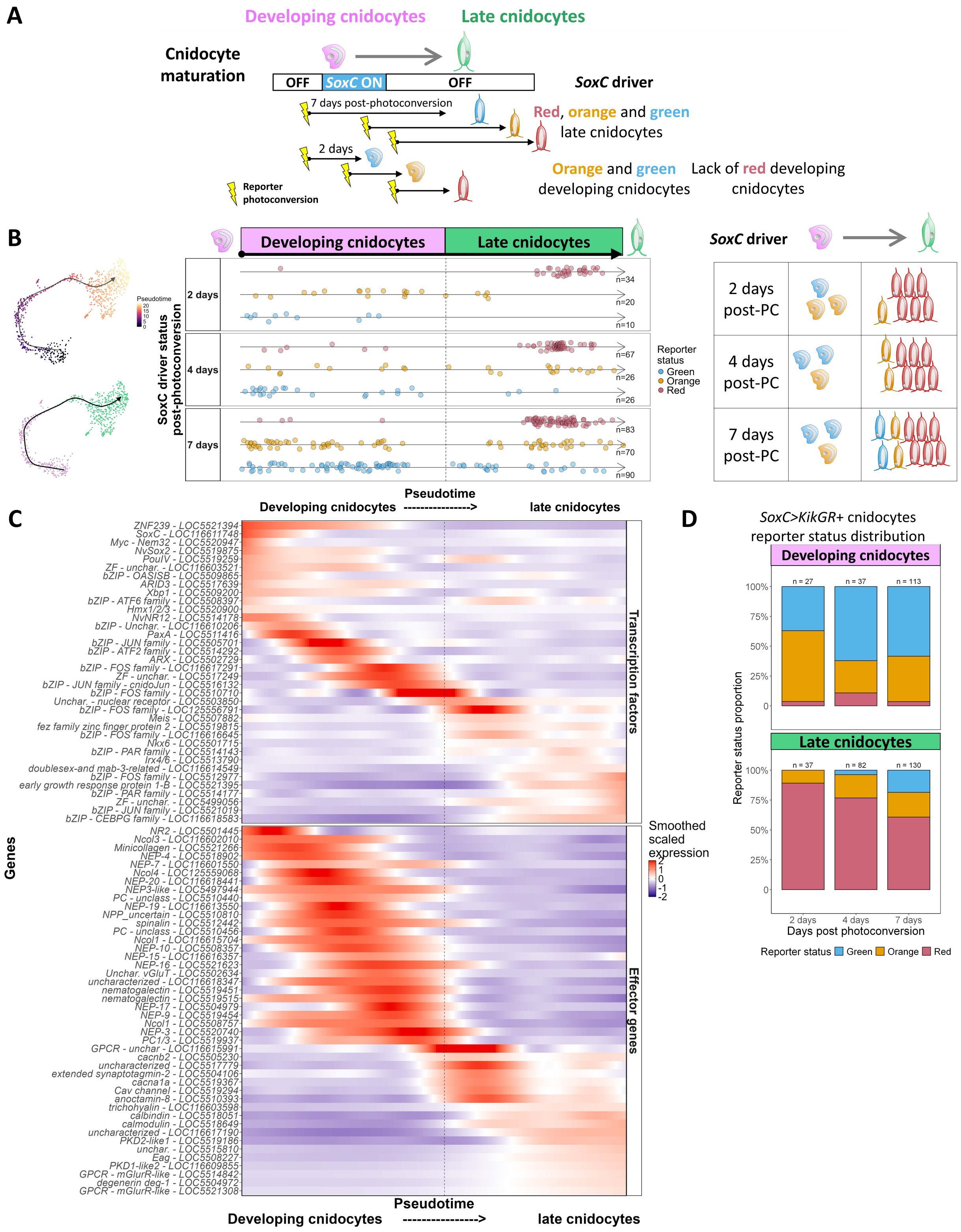

A
Developing cnidocytes
Late cnidocytes
Cnidocyte maturation
OFF
SoxC ON
OFF
SoxC driver
7 days post-photoconversion
Red, orange and green late cnidocytes
2 days
Orange and green developing cnidocytes
Lack of red developing cnidocytes
Reporter photoconversion
B
Pseudotime
Developing cnidocytes
Late cnidocytes
SoxC driver status post-photoconversion
2 days
4 days
7 days
n=34
n=20
n=10
n=67
n=26
n=26
n=83
n=70
n=90
Reporter status
Green
Orange
Red
Pseudotime
SoxC driver
2 days post-PC
4 days post-PC
7 days post-PC
C
Developing cnidocytes ----------------> late cnidocytes
Genes
ZNF239 - LOC5521394
SoxC - LOC116611748
Myc - Nem32 - LOC5520947
NvSox2 - LOC5519875
PouIV - LOC5519259
ZF - unchar. - LOC116603521
bZIP - OASISB - LOC5509865
ARID3 - LOC5517639
Xbp1 - LOC5509200
bZIP - ATF6 family - LOC5508397
Hmx1/2/3 - LOC5520900
NvNR12 - LOC5514178
bZIP - Unchar. - LOC116610206
PaxA - LOC5511416
bZIP - JUN family - LOC5505701
bZIP - ATF2 family - LOC5514292
ARX - LOC5502729
bZIP - FOS family - LOC116617291
ZF - unchar. - LOC5517249
bZIP - JUN family - cnidoJun - LOC5516132
bZIP - FOS family - LOC5510710
Unchar. - nuclear receptor - LOC5503850
bZIP - FOS family - LOC125556791
Meis - LOC5507882
fez family zinc finger protein 2 - LOC5519815
bZIP - FOS family - LOC116616645
Nkx6 - LOC5501715
bZIP - PAR family - LOC5514143
Irx4/6 - LOC5513790
doublesex-and mab-3-related - LOC116614549
bZIP - FOS family - LOC5512977
early growth response protein 1-B - LOC5521395
bZIP - PAR family - LOC5514177
ZF - unchar. - LOC5499056
bZIP - JUN family - LOC5521019
bZIP - CEBPG family - LOC116618583
NR2 - LOC5501445
Ncol3 - LOC116602010
Minicollagen - LOC5521266
NEP-4 - LOC5518902
NEP-7 - LOC116601550
Ncol4 - LOC125559068
NEP-20 - LOC116618441
NEP3-like - LOC5497944
PC - unclass - LOC5510440
NEP-19 - LOC116613550
NPP_uncertain - LOC5510810
spinalin - LOC5512442
PC - unclass - LOC5510456
Ncol1 - LOC116615704
NEP-10 - LOC5508357
NEP-15 - LOC116616357
NEP-16 - LOC5521623
Unchar. vGluT - LOC5502634
uncharacterized - LOC116618347
nematogalectin - LOC5519451
nematogalectin - LOC5519515
NEP-17 - LOC5504979
NEP-9 - LOC5519454
Ncol1 - LOC5508757
NEP-3 - LOC5520740
PC1/3 - LOC5519937
GPCR - unchar - LOC116615991
cacnb2 - LOC5505230
uncharacterized - LOC5517779
extended synaptotagmin-2 - LOC5504106
cacna1a - LOC5519367
Cav channel - LOC5519294
anoctamin-8 - LOC5510393
trichohyalin - LOC116603598
calbindin - LOC5518051
calmodulin - LOC5518649
uncharacterized - LOC116617190
PKD2-like1 - LOC5519186
unchar. - LOC5515810
Eag - LOC5508227
PKD1-like2 - LOC116609855
GPCR - mGlurR-like - LOC5514842
degenerin deg-1 - LOC5504972
GPCR - mGlurR-like - LOC5521308
Transcription factors
Effector genes
Smoothed scaled expression
2
1
0
-1
-2
Pseudotime
Developing cnidocytes ----------------> late cnidocytes
D
SoxC>KikGR+ cnidocytes reporter status distribution
Developing cnidocytes
n = 27
n = 37
n = 113
Late cnidocytes
n = 37
n = 82
n = 130
Reporter status proportion
100%
75%
50%
25%
0
2 days
4 days
7 days
Days post photoconversion
Reporter status
Green
Orange
Red

**Supplementary Figure 13: Biphasic transcriptional maturation of cnidocytes from *SoxC*+ cells with a primary capsule program followed by a secondary neural program delayed after capsule formation**

**A** Cnidocyte maturation is a two-step process with late cnidocytes (in green, represented in blue) arising from transiently expressing *SoxC*+ cells (capsule formation developing cnidocytes phase, in pink). **B** Cnidocyte temporal maturation. (**left**) All cnidocytes UMAP projection and slingshot-inferred pseudotime. (**middle**) Pseudotime alignment of *SoxC>KikGR*+ cnidocytes along their differentiation trajectory by reporter status (green, orange or red cells) at 2, 4 and 7 days post-photoconversion. Developing cnidocytes are green or orange at all timepoints post-photoconversion. A majority of non-expressing red mature cnidocytes are recovered, with an increasing fraction of newly generated orange and green late cnidocytes occurring at later days post-photoconversion. Vertical dashed line separates developing cnidocytes and late cnidocytes. (**right**) Temporal reconstruction of cnidocyte maturation from *SoxC>KikGR*+ labeled developing cnidocytes (as in Fig. 3A). **C** Transcription factors (TF) (**top**) and effector genes (**lower**) smoothed scaled expression across the pseudotime differentiation trajectory. Vertical dashed line separates developing cnidocytes and late cnidocytes which display distinct effector and transcription factor profiles. Transcription factors with known roles in capsule formation are expressed early, at the developing cnidocyte stage (*ZNF239, NvNR12, PaxA*), alongside the early *SoxC* expression. There is an abrupt shift to distinct transcription factors (several bZIP, *nkx6*, *FoxJ1*, a *Myc* and uncharacterized *Sox* gene) during maturation and expression of the secondary neural program. **D** Proportion of green, orange and red developing cnidocytes (**top**) and mature cnidocytes (**bottom**) at 2-, 4- and 7-days post *SoxC>KikGR*+ photoconversion, with the number of cells recovered indicated above. Same data as in panel B (middle). PC: Photoconversion.

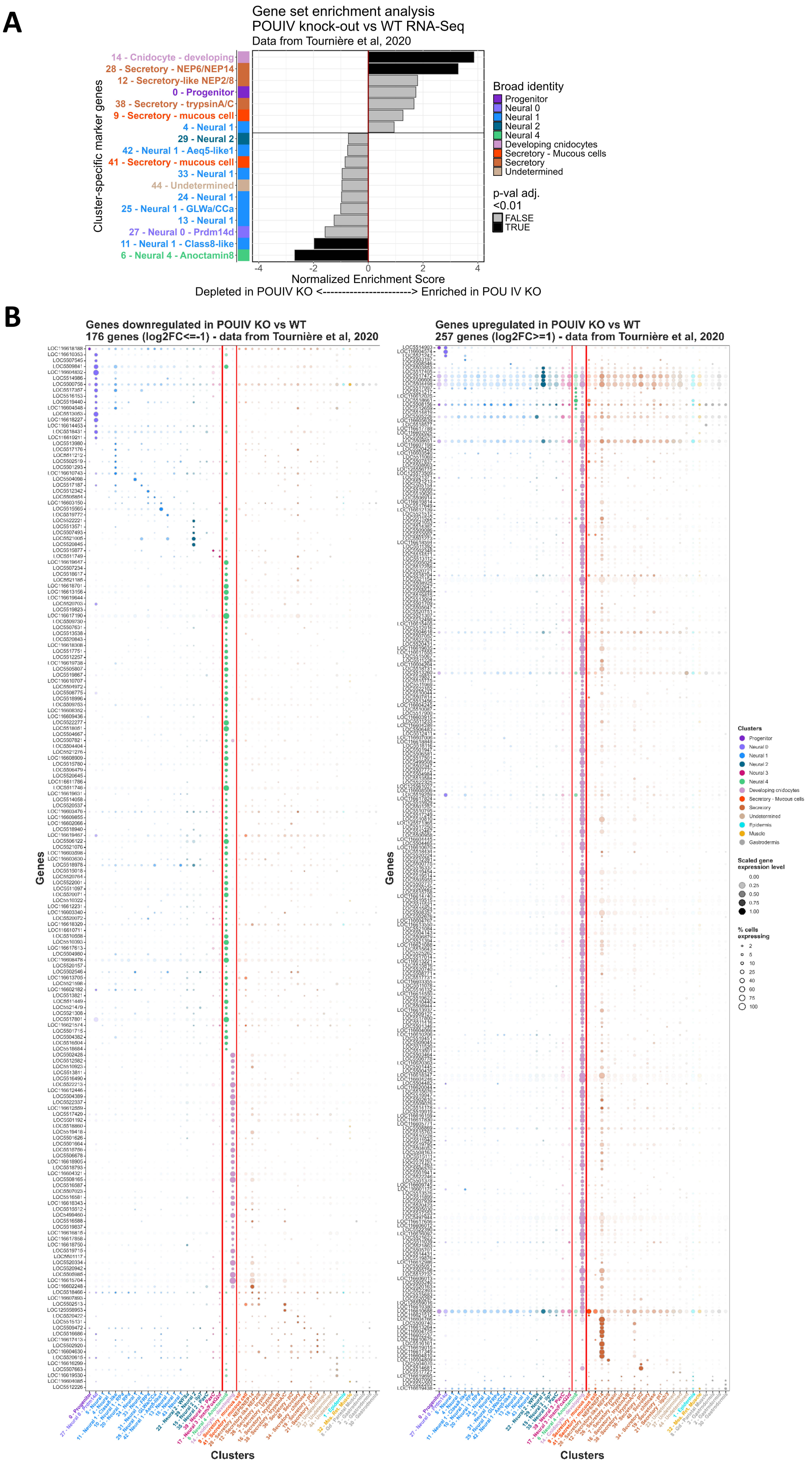
A
Gene set enrichment analysis
POUIV knock-out vs WT RNA-Seq
Data from Tournière et al, 2020
Cluster-specific marker genes
14 - Cnidocyte - developing
28 - Secretory - NEP6/NEP14
12 - Secretory-like NEP2/8
0 - Progenitor
38 - Secretory - trypsinA/C
9 - Secretory - mucous cell
4 - Neural 1
29 - Neural 2
42 - Neural 1 - Aeq5-like1
41 - Secretory - mucous cell
33 - Neural 1
44 - Undetermined
24 - Neural 1
25 - Neural 1 - GLWa/CCa
13 - Neural 1
27 - Neural 0 - Prdm14d
11 - Neural 1 - Class8-like
6 - Neural 4 - Anoctamin8
-4
-2
0
2
4
Normalized Enrichment Score
Depleted in POUIV KO <-----------------------> Enriched in POU IV KO
Broad identity
Progenitor
Neural 0
Neural 1
Neural 2
Neural 4
Developing cnidocytes
Secretory - Mucous cells
Secretory
Undetermined
p-val adj.
<0.01
FALSE
TRUE
B
Genes downregulated in POUIV KO vs WT
176 genes (log2FC<=-1) - data from Tournière et al, 2020
Genes upregulated in POUIV KO vs WT
257 genes (log2FC>=1) - data from Tournière et al, 2020
Genes
Clusters
Clusters
Progenitor
Neural 0
Neural 1
Neural 2
Neural 3
Neural 4
Developing cnidocytes
Secretory - Mucous cells
Secretory
Undetermined
Epidermis
Muscle
Gastrodermis
Scaled gene expression level
0.00
0.25
0.50
0.75
1.00
% cells expressing
2
5
10
25
40
60
75
100

**Supplementary Figure 14: Reanalysis of a *PouIV* KO dataset supports a biphasic transcriptional maturation of cnidocytes**

**A** Gene Set Enrichment Analysis (GSEA) showing the differential enrichment of cluster marker genes in *PouIV* KO RNA-Seq data [18]. Genes upregulated in the *PouIV* KO animals (Normalized Enrichment Score >0) significantly overlap with marker genes expressed in developing cnidocytes (in pink), and in the cnidocyte-associated secretory cell cluster as they also express cnidocyte genes. Genes downregulated in PouIV KO animals (Normalized Enrichment Score <0) significantly overlap with marker genes expressed in mature cnidocytes (in green, Neural 4 cell class). **B** Expression of top downregulated ($log2FC \leq -1$, **left**) and upregulated ($log2FC \geq 1$, **right**) genes in the *PouIV* KO RNA-Seq data across our global KikGR+ clustering (same cluster order and coloring as in Fig. 3G). Genes upregulated in *PouIV* KO animals are specifically expressed in mature cnidocytes (in green) and in developing cnidocytes (in pink), supporting a two-phase transcriptional program of cnidocyte maturation.

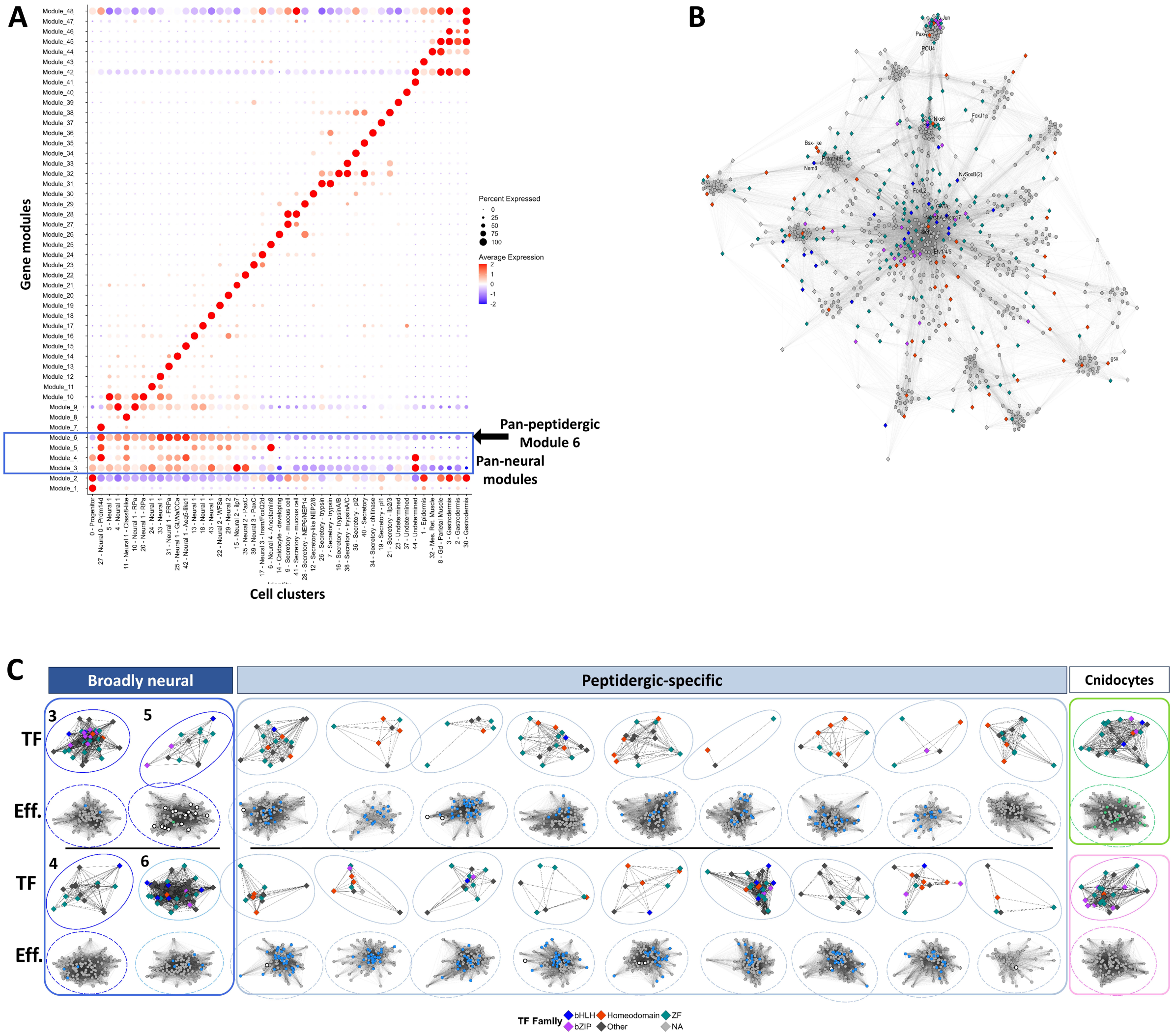


**Supplementary Figure 15: WGCNA gene modules**

**A** Module expression scores across clusters with pan-neural (Modules 3-6) and neural subtype-specific modules (Module 7-22 and 26). **B** Global neural modules co-expression network visualization (all transcription factors and top 20 genes in each module). **C** Neural WGCNA modules transcription factor (**top**) and top effectors (**bottom**) co-expression networks. Broadly neural modules (Modules 3-6) encompass bHLH (blue diamonds), bZIP (purple diamonds) and zinc finger (dark green diamonds) transcription factors. Peptidergic and late cnidocyte shared neural effector genes (in black in Fig. 6B) are in shared module 5 (in white circles). Each peptidergic subtype module (Modules 7-22) has homeodomain transcription factor(s) (red diamonds), zinc finger factors (dark green diamonds), and peptidergic-specific neural effector genes (blue circles, same genes as in Fig. 6B).

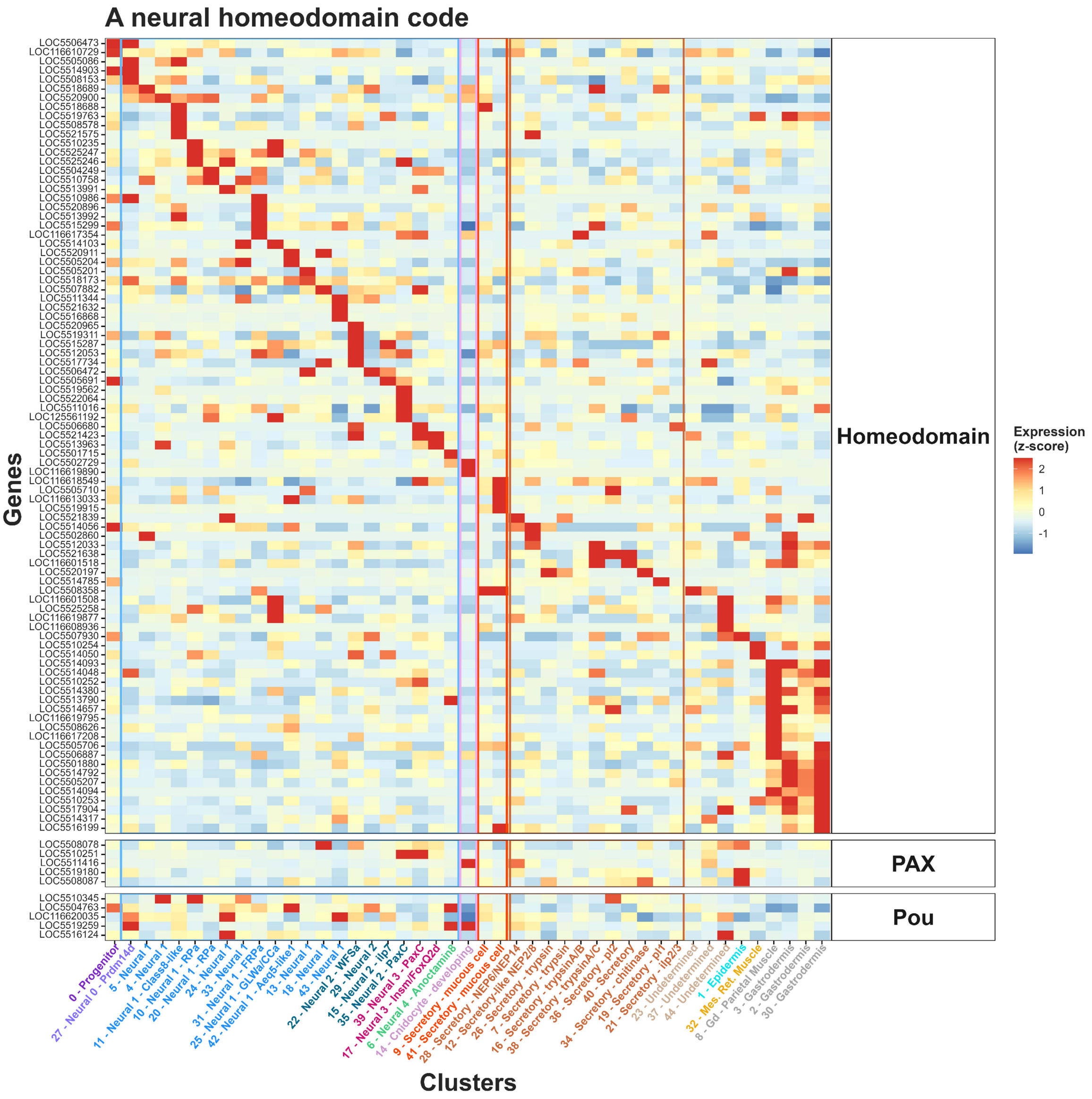


### Supplementary Figure 16: A neural homeodomain code

Expression of Homeodomain proteins across all clusters highlights neural subtype-specific expression, with each cluster expressing a unique set of transcription factors.
